# Soil microaggregate bacterial communities following *Amynthas tokioensis* and *Amynthas agrestis* earthworm co-invasion

**DOI:** 10.1101/2023.05.19.541528

**Authors:** Jaimie R. West, Bradley M. Herrick, Thea Whitman

## Abstract

Earthworms restructure the soil environment through burrowing, consumption, and casting behaviors. Though non-native European Lumbricid earthworms are well-studied in North American soils, the Asian pheretimoid *Amynthas tokioensis* and *Amynthas agrestis* earthworms exhibit distinct ecological patterns that alter invaded habitats. In particular, bioturbation may affect soil aggregation and microbial community assembly processes, such as dispersal and selection. We aimed to determine the effects of *A. tokioensis* and *A. agrestis* co-invasions in woodlands in Madison, WI, U.S. on soil bacterial communities and edaphic characteristics. Using 16S rRNA gene sequencing, we found that the presence and activity of these *Amynthas* species earthworms significantly affected bacterial community composition. At one site, there was a decrease in sample-to-sample dissimilarity (i.e., decreased beta diversity), with concomitant increases in homogenizing community assembly processes. However, at the other site, we found opposite trends, with evidence for increased compositional dissimilarity between samples and decreased evidence for homogenizing community assembly processes. Overall, inconclusive support for the hypothesized homogenization of bacterial community composition driven by homogenizing community assembly processes indicates that the effects of *Amynthas* pressure in these systems represent a departure from previously established soil disturbance paradigms. Instead, we conclude that aggregate formation via *A. tokioensis* and *A. agrestis* casting activity does not consistently impose a strong selective filter on soil bacterial communities, nor does the heightened earthworm activity necessarily act to meaningfully homogenize soil communities via dispersal. Overall increases in soil C and N under *Amynthas* spp. activity support previous work indicating enhanced decomposition and incorporation of soil litter, but future work could focus on long-term fate of microaggregate-protected C.

## 1. Introduction

Earthworm activity, soil aggregation and the soil microbial community interact in complex ways to influence the decomposition and storage of soil organic matter (SOM) and soil organic carbon (SOC). The specific effects of invasive European earthworms in the family Lumbricidae have been well-studied and documented in North American agricultural and forest soils (Burtelow et al., 1998; Hale et al., 2005; McLean et al., 2006; Yavitt et al., 2015). However, a “second wave” of invasion by ecologically distinct Asian pheretimoid earthworms in the family Megascolecidae is further altering the soil environment in woodland and urban natural areas throughout the eastern and central U.S., and portions of southeastern Canada (Chang et al., 2021). *Amynthas tokioensis* and *Amynthas agrestis* “jumping worms” are of particular concern because they spread quickly, eat voraciously to support rapid growth within an annual life cycle, and live at high densities, collectively producing a distinctive granular casting layer on the soil surface. A thorough review by Chang et al. (2021) details the biology, invasion history, and known effects of these earthworms, and also frames key questions and knowledge gaps that remain, which include the ecological impacts of *A. tokioensis* and *A. agrestis* regarding soil nutrient dynamics and the soil microbial community. The ecology and feeding habits of these *Amynthas* spp. earthworms and their effects on edaphic properties, soil structure, and the soil microbial community are species-specific (Chang et al., 2016a; Price-Christenson et al., 2020; Richardson et al., 2022), though specific effects are often obscured given that *Amynthas* typically co-occur in assemblages of two or three species (Chang et al., 2018). Results from the field have contradicted short-term laboratory mesocosm experiments (Chang et al., 2017; Greiner et al., 2012; Richardson et al., 2022), indicating that natural soils behave differently than highly disturbed mesocosms, while identifying a need for more field experiments (Chang et al., 2021).

First noted in Wisconsin at the University of Wisconsin (UW)–Madison Arboretum in Madison, WI, U.S. in 2013, *A. tokioensis* and *A. agrestis* earthworms primarily live at the soil-litter interface (“epi-endogeic”), feeding on both SOM and leaf litter, and produce castings on the soil surface rather than burrowing into mineral horizons. These *Amynthas* spp. grow and spread quickly due to dietary flexibility and aptitude for outcompeting existing earthworm populations (Chang et al., 2016b; Zhang et al., 2010), increasing their coverage by over 70% in one year at the UW Arboretum (Laushman et al., 2018). The co-invasion by *A. tokioensis and A. agrestis* has been associated with reduced leaf litter at the UW–Madison Arboretum (Qiu and Turner, 2017), potentially increasing primary productivity by mineralizing litter-derived nutrients (Laushman et al., 2018), though this increase in plant-available nutrients, particularly at the soil surface, is susceptible to loss via erosion, leaching or volatilization (Price-Christenson et al., 2020; Qiu and Turner, 2017; Yavitt et al., 2015). On the other hand, a short-term incubation experiment suggested that the encapsulation of litter and SOM via accelerated and enhanced biogenic aggregation driven by *A. agrestis* may increase SOM persistence (Zhang et al., 2013).

The drilosphere encompasses earthworm-affected soil, including casts and burrows, and can act as a hotspot of microbial activity and resource availability due to enhanced lability of ingested and excreted litter, secretion of earthworm mucus, and translocation of resources and microbes via earthworm cast deposition (Medina-Sauza et al., 2019). However, variations in earthworm ecology and diet drive nuanced effects on microbial communities and microbially-mediated SOM and SOC transformations. For example, litter-feeding may increase microbial biomass by enhancing nutrient availability and microbial niche partitioning (Chang et al., 2017), but SOM-feeding may suppress microbial biomass by reducing resource availability or through direct consumption of microbes (Chang et al., 2016a; Drake and Horn, 2007). The gut microbial community reflects the microbiome of the ingested material, with quantitatively insignificant abundances of endemic gut microbes (Drake and Horn, 2007). Thus, different earthworm species may have varying effects on the soil microbial community due to different dietary preferences. *A. tokioensis* and *A. agrestis*, which feed on both litter and SOM, harbor distinct gut microbial assemblages, which are apparent in fresh casts; however, the soil microbial community is not dominated by prevalent worm gut or cast taxa of either species (Price-Christenson et al., 2020). Passage through the anoxic, nutrient-rich earthworm gut alters microbial composition and activity (Drake and Horn, 2007), albeit for perhaps a relatively brief duration (McLean et al., 2006). The resulting microbial community is then subject to anoxic microsites created in dense, low-porosity worm castings (Bossuyt et al., 2005; Mummey et al., 2006).

Applying high-throughput sequencing to soil microaggregate fractions under *Amynthas* pressure will allow us to better understand how worm-soil-microbe interactions shape bacterial community composition and soil biogeochemistry. While certain trends exist in the literature regarding microbial communities of microaggregate fractions (e.g., Trivedi et al., 2017; Bach et al., 2018), these findings may not apply to earthworm-generated biogenic aggregates, which are formed via distinct processes and thus likely demonstrate unique microbial community composition patterns (Bossuyt et al., 2004; Mummey et al., 2006; Yavitt et al., 2015). That said, aggregate size fractions are operationally defined by resistance to stress, and are indicative of protective mechanisms and stability, regardless of origin. It follows that the typical aggregate-associated physical and chemical protections to SOM and SOC apply to earthworm-generated aggregates (Zhang et al., 2013). Macroaggregates (250-2000 µm) are relatively weakly woven together by roots and hyphae, whereas microaggregates (53-250 µm) are glued by physicochemical interactions and biomolecules, offering longer turnover times (Davinic et al., 2012; DeGryze et al., 2006; Totsche et al., 2018). Though earthworm digestion disrupts preexisting microaggregates, comminuted litter fragments—thoroughly mixed and encrusted with clay, microbiota, and mucus—are packed into castings where the organic fragments serve as nuclei for new aggregates (Barois et al., 1993; Bossuyt et al., 2006; Shipitalo and Protz, 1989).

Whereas the process of microaggregate formation may gradually occur via microbial processing of organic matter within macroaggregates (Six et al., 1999) and physical processes such as wet-dry cycles (Dexter et al., 1988), microaggregate formation via earthworm processing is relatively rapid; one study demonstrated that earthworm activity resulted in enrichment of ^13^C-labelled litter in occluded microaggregates after twelve days, whereas in the absence of earthworms, labelled litter was incorporated into macroaggregates (outside of occluded microaggregates) (Bossuyt et al., 2004). This has important implications because macroaggregates have shorter turnover times compared to microaggregates, and thus SOM contained in macroaggregates is more vulnerable to microbial decomposition, whereas the occluded microaggregate-SOM remains protected following macroaggregate turnover (Bossuyt et al., 2006). Further, earthworm cast microaggregates offer an increased level of physical protection due to high stability imparted through thixotropic hardening/aging, which strengthens the organo-mineral associations (Shipitalo and Protz, 1989), while engendering poor conditions for microbial activity (e.g., low porosity for microbial habitat and gas/water exchange) (Jouquet et al., 2008; Lavelle et al., 2020). These distinctions in microaggregate formation and microbial habitat impact community compositions: whole microaggregate communities vs. interior microaggregate communities have been found to be similar for microaggregates created by earthworms, whereas gradually-formed microaggregates host distinct bacterial community compositional profiles on their interior, indicating habitat niche development (Mummey et al., 2006).

We were first drawn to this earthworm invasion as a modality of intrinsic soil mixing in natural soils (sensu Rillig et al., 2016). West and Whitman (2022) showed that repeated soil mixing disturbances in the lab reduced bacterial richness, while favoring organisms with potential for faster growth rates. These findings were echoed under tillage disturbance (West et al., 2023), even though tillage occurred much less frequently relative to the frequent disturbance that was applied in the lab experiment. Because *A*. *tokioensis and A*. *agrestis* live in high density populations and are restricted to the soil surface, we anticipated findings would reflect the lab mixing experiment, with strong indicators of frequent and thorough soil mixing. Given the rapid encapsulation of ingested soil, litter, and microbial communities via earthworm activity, the effects on microbial diversity and ecology are unclear (Bossuyt et al., 2005; Mummey et al., 2006). Highly stochastic assemblages of microbial communities may persist, given suppressed microbial activity in the well-protected and potentially anoxic cast microenvironment (Bossuyt et al., 2005; Jouquet et al., 2008; Mummey et al., 2006), as compared to more selectively assembled communities under microbially-mediated aggregate formation.

Community assembly processes influence community composition to generate ecological relationships. The community assembly processes (Vellend, 2010) of interest in this study are as follows: Dispersal describes the generally stochastic movement and establishment of organisms, and may occur in soil via physical disturbance or pore water movement (Zhou and Ning, 2017). Homogenizing dispersal increases compositional similarity between communities, whereas dispersal limitation increases compositional differences between communities, potentially allowing for stochastic demographic changes to community composition — termed ‘drift’ (Stegen et al., 2013). Selection refers to deterministic or niche-based processes dictated by biotic factors, such as inter-taxa fitness differences, and abiotic factors, such as environmental filters (Hutchinson, 1957). Homogeneous selection describes increased phylogenetic similarity between communities attributable to similar habitats or filters (Dini-Andreote et al., 2015). Variable selection decreases phylogenetic similarity between communities due to variable conditions (Stegen et al., 2015). The relative influences of these community assembly processes can be statistically inferred for separate ‘bins’ of phylogenetically-related OTUs, thus enabling a collective representation of the various community assembly processes that may influence subsets of community members (Ning et al., 2020). This method compares observed phylogenetic distance and compositional dissimilarity metrics between communities to null models of stochastically assembled communities (Stegen et al., 2015, 2013, 2012).

We sought to better understand how bacterial communities of surface soil and its microaggregate fractions are affected by co-invasion by *Amynthas tokioensis* and *A. agrestis*, and how these effects are modulated through changes to soil aggregation. Better understanding the effects of *Amynthas* spp. invasion on microbial community composition and community assembly in microaggregate environments can improve understanding of SOM protection and SOC persistence, with findings also applicable to soils affected by other earthworms. We collected soil samples from invaded and uninvaded areas of two woodlands in Madison, WI, U.S., and analyzed edaphic properties, bacterial community composition, diversity, and community assembly processes of the bulk soil, free microaggregate, and occluded microaggregate fractions, using 16S rRNA gene amplicon sequencing. We hypothesized:

H1) Under *Amynthas* spp. pressure, there would be increased sample-to-sample similarity in community composition (i.e., lower beta diversity), and community assembly would be influenced by homogenizing dispersal and homogeneous selection, whereas in the control, community assembly would be influenced by dispersal limitation and variable selection.

H2) Under *Amynthas* spp. pressure, the free and occluded microaggregate fractions would support similar bacterial communities, whereas in the control, the free and occluded microaggregate fractions would support bacterial communities that are more distinct from each other. This hypothesis is built on the idea that *A. tokioensis* and *A. agrestis*-driven aggregate formation rapidly occludes free microaggregates into macroaggregate structures, rather than more gradual occluded microaggregate formation within macroaggregates.

## 2. Methods

### 2.1 Soil collection

Soil was sampled from two separate woodland areas in Madison, WI, U.S.: 1) the UW–Madison Arboretum in the Gallistel Woods management unit (hereafter: Gallistel) (43°2’38”N, 89°25’25”W, 263 m a.s.l.) on Virgil silt loam (fine-silty, mixed, superactive, mesic Udollic Endoaqualfs), Dodge silt loam (Fine-silty, mixed, superactive, mesic Typic Hapludalfs), and Wacousta silty loam (Fine-silty, mixed, superactive, mesic, Typic Endoaquolls) soils in an *Acer saccharum*-dominated mesic forest with *Quercus alba* and *Tilia americana* (partially cleared but never plowed); and, 2) the UW–Madison Lakeshore Nature Preserve (hereafter: Lakeshore) in Wally Bauman Woods (43°5’17”N, 89°26’32”W, 275 m a.s.l.) and Frautschi Point Woods (43°5’31”N, 89°25’54”W, 268 m a.s.l.) on Whalan, McHenry, and Dodge silt loam soils (Fine-loamy and fine-silty, mixed, superactive, mesic Typic Hapludalfs) under a canopy dominated by *Acer saccharum* with *Tilia americana* and *Prunus serotina* (Fig. S1).

These areas are under co-invasion by *A. tokioensis* and *A. agrestis* (Price-Christenson et al., 2020; Richardson et al., 2022); we will collectively refer to this co-invasion as “*Amynthas* pressure”, while noting that not all earthworms of the genus *Amynthas* share the same ecological traits. Gallistel samples were collected from high *Amynthas* pressure and low *Amynthas* pressure plots, which were chosen based on worm surveys conducted 2015–2018, and 2021; plot establishment and data from 2015 and 2016 were detailed and reported in Laushman et al. (2018). Control plots were also sampled from an adjacent area of Gallistel that had not been invaded. Lakeshore samples were collected from high *Amynthas* pressure plots in Wally Bauman Woods, based on worm survey data collected in 2017, 2018, and 2021 (data not yet reported), and uninvaded control plots in Frautschi Point Woods, established and surveyed in 2021. High *Amynthas* pressure plots had relatively abundant worm counts in the first survey year (2015 or 2017, depending on site), potentially indicating a longer time since invasion, and a consistently high population in each year surveyed (mean 18 worms/0.36 m^2^ plot/year, range of 10-35 worms), whereas low *Amynthas* pressure plots were more recently invaded, with the first *Amynthas* worms noted in 2018, and no particularly abundant years (mean 7 worms/0.36 m^2^ plot/year, range of 2-12 worms). Three plots were sampled for each treatment (30 October, 2021 at Gallistel and 13 November, 2021 at Lakeshore; Fig. S1), collecting five intact cores at least 0.5 m beyond the perimeter of the permanent plots, for a total of 15 cores per treatment.

Due to our interest in discerning dispersal processes, our sampling design focused on ensuring relatively high spatial proximity of individual cores within a given plot. Cores were 7.9 cm dia., evenly spaced just within the perimeter of a 48 cm dia. circle; distance between adjacent cores was approximately 15 cm. As detailed below, the top 3 cm were analyzed in order to target soil where these *Amynthas* spp. are most active (Qiu and Turner, 2017; Richardson et al., 2009). Intact cores were temporarily kept in a cooler, and then held at 4 °C for up to ten days until sample processing.

### 2.2 Sample processing and aggregate size fractionation

In order to assess variability in community composition and community assembly at a relatively small spatial scale, each core was processed separately (Fig. 1). The top 3 cm of each field-moist soil core were gently passed through a 2 mm sieve (henceforth referred to as “bulk” soil). Then, 80 g of this field-moist bulk soil was subjected to aggregate size fractionation via wet sieving (Elliott, 1986) to isolate the macroaggregate fraction (250 µm–2000 µm), free microaggregate fraction (53 µm–250 µm), and the silt + clay-sized fraction (< 53 µm). Then, 20 g of the moist macroaggregate fraction was separated into occluded fractions via rapid shaking with glass beads to break up the macroaggregates under water flow to encourage liberated microaggregates and other material to pass through a 250 µm sieve, as previously described (Six et al., 2002, 2000); macroaggregate-occluded fractions included the occluded microaggregate fraction (53 µm–250 µm), occluded silt + clay-sized fraction (< 53 µm), and occluded coarse POM + coarse sand-sized fraction (250 µm–2000 µm). Other modifications to the wet sieving method included a slaking for two minutes prior to the first wet sieving step and draining each wet sieved fraction for two minutes prior to subsampling as described below. The primary objective of fractionation was to isolate the free and occluded microaggregate fractions for community analysis, but the relative dry mass of each size fraction was also determined. Correction for sand content of aggregate fractions was not performed and thus all size fractions also consisted of the primary mineral particles of that size.

**Figure 1.**
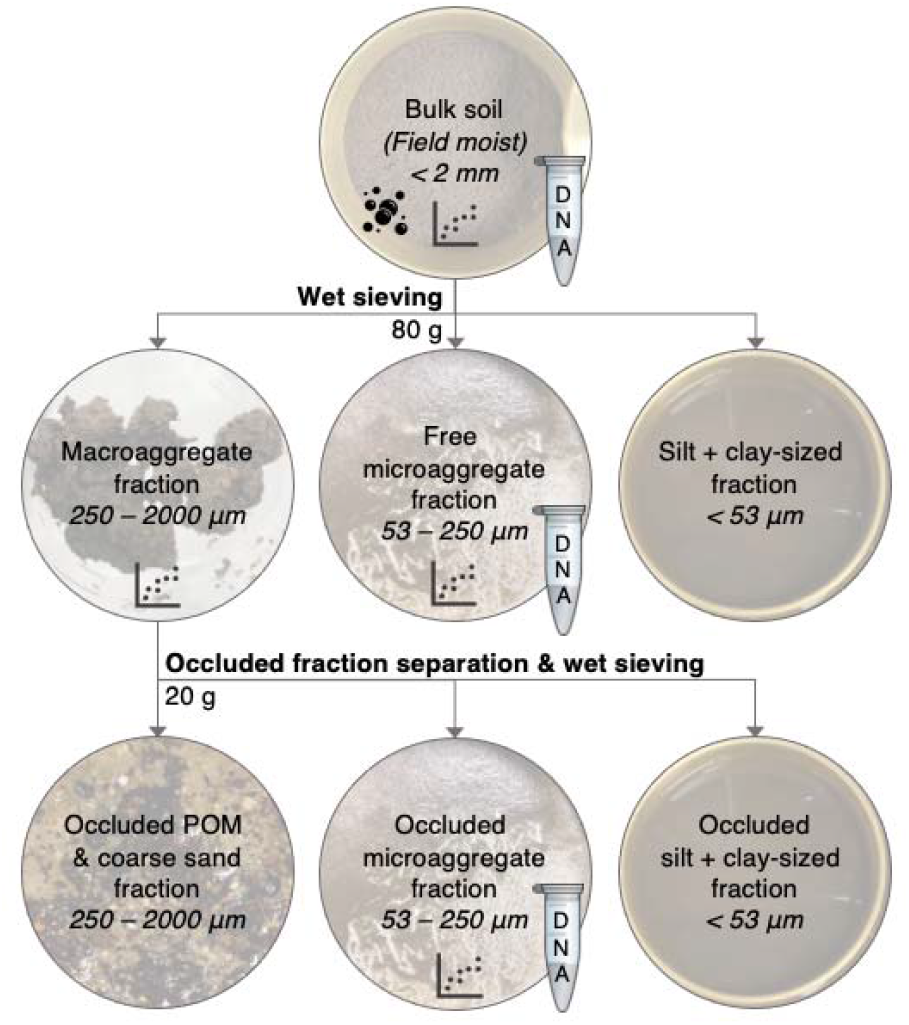
Aggregate size fractionation schematic. Bulk soil (80 g) was subjected to wet sieving to separate macroaggregate (250*–*2000 µm), free microaggregate (53–250 µm), and silt + clay-sized (< 53 µm) fractions. A 20 g (wet) subsample of the macroaggregate fraction was then further separated into occluded microaggregate (53–250 µm), occluded silt + clay-sized (< 50 µm), and occluded POM + coarse sand fractions (250–2000 µm). The “DNA” tube indicates that subsamples were retained for 16S rRNA gene amplicon sequencing. The graph icon indicates that subsamples were collected to measure total carbon and total nitrogen. The bubble icon indicates that soil respiration was measured on bulk soil.

Moisture content was estimated for bulk soil, the macroaggregate fraction, the free microaggregate fraction, and the occluded microaggregate fraction by drying subsamples in a 60 °C oven for 24 hours. Bulk, free microaggregate, and occluded microaggregate soil was subsampled for DNA extraction (see section 2.5), and bulk soil was also subsampled to measure soil respiration (see section 2.3). The remaining wet-sieved soil was washed from each sieve (or washbasin) into aluminum pans to determine the dry mass of each fraction. Overall recovery (macroaggregate + free microaggregate + [silt + clay] fractions) was 97–100% for all treatments at both sites, and macroaggregate recovery (occluded microaggregate + [occluded silt + clay] + occluded coarse POM) was 97–101% for all treatments.

### 2.3 Soil analysis

The bulk soil (sieved to < 2 mm), macroaggregate, free microaggregate, and occluded microaggregate fraction subsamples that were retained for dry mass conversion were ground to a powder and used to quantify total soil carbon and nitrogen by flash combustion with a Flash EA 1112 CHN Automatic Elemental Analyzer (Thermo Finnigan, Milan, Italy) and soil pH (excluding the macroaggregate fraction). Soil pH was determined from the supernatant of a 1:1 soil:CaCl_2_ (0.1 M) slurry using a micro pH electrode (modified from Braus and Whitman, 2021). For routine soil analysis, a composite sample representing each treatment was comprised of an equal mass of bulk soil from each plot, and submitted to the UW Soil and Forage Analysis Lab (Madison, WI, U.S.) to determine soil texture, organic matter content, pH, and plant-available P, K, Ca and Mg. Routine soil properties are reported in Table S1.

Soil respiration (CO_2_ evolution) from sieved fresh soil was measured using the MicroResp system (James Hutton Ltd., Aberdeen, Scotland) following general instructions for use (Campbell et al., 2003) without added substrate. At the time of aggregate fractionation, 300 mg of freshly sieved (< 2 mm), field-moist soil from each soil core was placed into each of six wells of a deep-well plate. Wells were covered and stored at 4 °C for up to six hours. Each deep-well plate, containing up to ten different samples, was covered in parafilm and firmly tapped on the benchtop 20 times to minimize large air pockets. The deep-well plate was then incubated at 25 °C in a CO_2_-free environment overnight (approximately 16 hours), to help deplete CO_2_ from the well headspace and soil air. The following day, a colorimetric detection plate was read at absorbance wavelength 570 nm using a BioTek Synergy 2 spectrophotometer microplate reader (Agilent, Santa Clara, CA). After confirming that all wells had highly similar readings (< 5% coefficient of variance), the detector plate was inverted over the deep-well plate, connected by the 96-well seal, and clamped together. After six hours of incubation at 25 °C, the clamp set-up was dismantled, and the colorimetric plate was immediately re-read to determine CO_2_ evolution. The colorimetric agar in the detection plates was made per the MicroResp manual instructions (version 4), and CO_2_ evolution was calculated following the manual’s instructions, with a minor modification to subtract CO_2_ values for a set of empty deep wells in each plate (blanks).

### 2.4 Worm cast collection

To collect *Amynthas* worm casts for microbial community analysis, earthworms were collected on 18 October, 2021 by pouring a solution of 20 g mustard powder shaken and suspended in 2 L of water over an area of 0.23 m^2^ near each plot (Gunn, 1992; Laushman et al., 2018). Earthworms that emerged were rinsed in sterile 0.85% NaCl solution and individually placed in sterile petri dishes with a moist paper towel. Petri dishes were held under dark, room temperature conditions until each worm produced castings (max. 24 hrs). Worm castings were composited by plot and subjected to DNA extraction and 16S rRNA gene sequencing as detailed below. Worms were all visually identified in the field as belonging to the genus *Amynthas*. Worm surveys conducted in late summer 2021 identified 87% of the worms in the Gallistel plots as *A. tokioensis*, and the remainder as *A. agrestis*, and 100% of the worms in the Lakeshore plots as *A. tokioensis*. Shallow mustard pours were also performed near control plots to confirm that no *Amynthas* were present; there were also no other worms present in the control plots, though, the volume of mustard solution applied would not necessarily have brought anecic earthworms to the surface.

### 2.5 DNA extraction and 16S rRNA gene sequencing

Total genomic DNA was extracted from bulk soil, free microaggregate, and occluded microaggregate soil fractions using the DNeasy PowerLyzer PowerSoil Kit (Catalog No. 12855, Qiagen, Germantown, MD), following manufacturer’s instructions. We used 250 mg samples of field-moist bulk soil for DNA extraction, but, due to the wetness of the microaggregate fractions following wet sieving, we used 450 mg samples of these fractions in order to approximate the same dry-mass equivalent of 250 mg of field-moist bulk soil, based on preliminary testing. The microaggregate samples were transferred directly from the drained sieves into the DNA extraction tubes, which were immediately frozen at −20 °C, and stored at −80 °C for up to three months prior to DNA extraction. Complete library preparation details can be found in the Supplementary Information. Briefly, the 16S rRNA genes of extracted DNA were amplified in triplicate using PCR. Variable region V4 of the 16S rRNA gene was targeted using forward primer 515f and reverse primer 806r (Walters et al., 2016). Primers also contained barcodes and Illumina sequencing adapters (Kozich et al., 2013). The following reagents comprised each 25 μL PCR reaction: 12.5 μL Q5 Hot Start High-Fidelity 2X Master mix (Catalog No. M0494, New England BioLabs, Ipswich, MA), 1.25 μL 515f forward primer (10 mM), 1.25 μL 806r reverse primer (10 mM), 1 μL DNA extract, 1.25 μL Bovine Serum Albumin (20 mg/mL; Catalog No. 97064-342, VWR International, Radnor, PA), and 7.75 μL PCR-grade water. The plate was sealed and briefly centrifuged prior to 30 PCR cycles on an Eppendorf Mastercycler nexus gradient thermal cycler (Hamburg, Germany) using the following parameters: 98 ^∘^C for 2 min + 30 × (98 ^∘^C for 10 seconds + 58 ^∘^C for 15 seconds + 72 ^∘^C for 10 seconds) + 72 ^∘^C for 2 min and 4 ^∘^C hold. Amplified DNA was confirmed via gel electrophoresis, then normalized and purified, prior to paired-end 250 base pair sequencing on an Illumina MiSeq sequencer at the UW–Madison Biotech Center. To obtain high coverage, the same library was sequenced twice under identical conditions, and total reads were pooled for each sample after processing as described next. Sequencing data was processed using a QIIME2 (Bolyen et al., 2019) pipeline, with DADA2 (Callahan et al., 2016) as the operational taxonomic unit (OTU, or amplicon sequence variant)-picking algorithm, and taxonomy assignment using the SILVA 132 reference database (Quast et al., 2013; Yilmaz et al., 2013). This yielded 12,957,889 demultiplexed sequences, which was reduced to 8,063,244 after denoising, with a mean length of 227 base pairs (SD = 2.2). Excluding extraction blanks, a total of 21,531 OTUs were identified. Amplicon sequences [will be] available in the National Center for Biotechnology Information (NCBI) Sequence Read Archive (SRA) under accession [TBD]. Our primers targeted both bacteria and archaea, but because our communities were dominated by bacteria (98.4% of total reads), for simplicity, we will refer to bacteria when discussing communities in this manuscript. Over 98% of archaeal reads represented the phylum *Crenarchaeota*.

### 2.6 Data analysis

Data analysis was performed in R (R-Core-Team, 2018), using *ggplot2* (Wickham, 2016) for data visualization. The R code used to perform these analyses and to create the following figures is available at https://github.com/jaimiewest/Soil-Disturbance-Amynthas. To test for a significant effect of *Amynthas* pressure on proportion of soil in each fraction, C content of each fraction, and respiration, we used ANOVA followed by Tukey’s HSD post-hoc comparison for significant results (*p* < 0.05). To test for a significant effect of *Amynthas* pressure, soil fraction, or interaction of these factors on soil C content, soil N content, and soil C:N ratio, we performed ANOVA as described above. Unless otherwise noted, reported *p* values refer to ANOVA tests. Community composition was visualized using principal coordinates analysis (PCoA) created with the *ordinate* function in the *phyloseq* package for R (*phyloseq::ordinate*) (McMurdie and Holmes, 2013) using Bray-Curtis dissimilarities (Bray and Curtis, 1957) of Hellinger-transformed relative abundance data (Legendre and Gallagher, 2001). To test for a significant effect of *Amynthas* pressure, soil fraction, or interaction of these factors on community composition, we used permutational multivariate analysis of variance (PERMANOVA) to partition Bray-Curtis dissimilarity matrices among sources of variation using *vegan::adonis2* (Anderson, 2001). A significant result (*p* < 0.05) was subjected to *post-hoc* pairwise factor comparisons, adjusting p-values using the Benjamini-Hochberg method (Benjamini and Hochberg, 1995) to identify significant differences. To compare differences in community composition due to *Amynthas* pressure or soil fractions, we tested for homogeneity of multivariate dispersions (PERMDISP; *vegan::betadisper*) (Anderson, 2006), using ANOVA to test the distances to group spatial median. Further, we also evaluated the effect of *Amynthas* pressure on dispersion of free and occluded microaggregate fraction communities within each soil core. To describe richness, we used the weighted linear regression model of OTU richness estimates, which weights observations based on variance, using *breakaway::betta* (Willis and Bunge, 2014). We also calculated Faith’s phylogenetic diversity (PD) (Faith, 1992; Pérez-Valera et al., 2015) to assess differences in phylogenetic distance (i.e., sample branch length) using *picante::pd* (Kembel et al., 2010).

To further understand changes in community composition, we calculated the weighted mean predicted 16S rRNA gene copy number (Nemergut et al., 2016), which has been shown to correlate with potential growth rate (Klappenbach et al., 2000) and disturbance (West et al., 2023; West and Whitman, 2022; Whitman et al., 2019), and compared treatments and fractions using ANOVA and *post-hoc* testing, as described above. 16S rRNA gene copy numbers were predicted using the ribosomal RNA operon database (rrnDB) (Stoddard et al., 2015).

After evaluating our key questions, we used differential abundance to identify significant treatment-driven shifts in relative abundances of individual taxa. For this analysis, we compared the *Amynthas* pressure treatments to the control (excluding taxa with mean relative abundance < 0.00001) and subjected these data to a beta-binomial regression model and “Wald” hypothesis test in *corncob*∷*differentialTest* (Martin et al., 2021), which controls for the effect of the treatment on dispersion. We report the µ value, which is the coefficient used to estimate relative abundance in the *corncob* model and is proportional to the fold-change in relative abundance between the treatment and control. We also assessed differential abundances of taxa in the microaggregate fractions as compared to the bulk soil communities.

### 2.7 Community assembly process assignment

In order to determine the influential community assembly processes within each treatment and fraction, we first applied a method developed by Stegen et al. (2012, 2013, 2015), which compares sample pairs of interest (i.e., each possible pair of samples from the same site, *Amynthas* pressure treatment, and fraction) to null models constructed using the full dataset that represent stochastic assembly, in order to determine the relative influence of selection (based on phylogenetic distances), or dispersal (based on compositional dissimilarities), as detailed in the Stegen et al. papers. Briefly, the influence of selection was first tested using the abundance-weighted beta-mean nearest taxon distance (βMNTD; the mean phylogenetic distance between each OTU in one community and its closest relative in another community) (*picante::comdistnt*) (Kembel et al., 2010). Homogeneous selection was identified in comparisons for which βMNTD was more than 2 standard deviations below the mean of the null distribution, indicating lower mean phylogenetic distance between pairwise communities than observed in the null. Variable selection was identified in comparisons for which βMNTD was more than 2 standard deviations above the mean of the null distribution, indicating higher mean phylogenetic distance between pairwise communities than observed in the null. Comparisons that fell within 2 standard deviations of the null mean were considered to lack a dominant influence of selection, and were subsequently tested for the influence of dispersal using the modified Raup-Crick metric based on Bray–Curtis dissimilarities (RC_Bray_) (*phyloseq::distance*) (Chase et al., 2011). Homogenizing dispersal was identified in comparisons for which RC_Bray_ was significantly lower than the mean of the null distribution, indicating a higher level of similarity between community compositions than was observed in the null condition; and dispersal limitation was identified in comparisons for which RC_Bray_ was significantly higher than the null mean, indicating lower similarity. Comparisons that were similar to the null mean for both metrics were considered undominated by any particular community assembly process.

In addition to determining the dominant community assembly processes at the full-community scale, we also assigned community assembly processes at a finer taxonomic scale, by separately assessing phylogenetically-related bins of OTUs (*iCAMP::pdist.big* and *iCAMP::icamp.big*), as detailed by Ning et al. (2020). This approach captures the various ecological mechanisms governing community assembly of the microbial subcommunities within each soil sample. Like the above-detailed full community assessment, here we also compared βMNTD and RC_Bray_ metrics to null models in order to determine the influences of selection and dispersal, respectively, though sample-to-sample comparisons were made within phylogenetic bins, and the dominant process was weighted by that bin’s relative abundance. We used the default parameters as detailed in Ning et al. (2020) and the R documentation (i.e., minimum of 24 OTUs per bin, confirmed by phylogenetic signal testing using *iCAMP::dniche* and *iCAMP::ps.bin*; phylogenetic null model randomization within bins; taxonomic null model randomization across all bins), with the exception of the phylogenetic distance metric, for which we used βMNTD to better compare results with the full-community scale assessment. To test for a significant effect of *Amynthas* pressure treatment on the influence of community assembly processes that had > 5% influence, we performed ANOVA as described above. Community relative abundance data was Hellinger-transformed (Legendre and Gallagher, 2001) for both community assembly assessments.

## 3. Results

### 3.1 *Amynthas* pressure increased carbon in macroaggregate and occluded microaggregate fractions

High *Amynthas* pressure increased total soil carbon content at Lakeshore by 50% (Fig. 2B, *p* < 0.001), and at Gallistel by 19% (Fig. 2A, trend not significant), reported here on a per unit of bulk soil basis. Carbon content of the macroaggregate and occluded microaggregate fractions were also significantly higher under high *Amynthas* pressure at both sites (Fig. 2A and B, *p* < 0.01). Of the increase in total soil C at each site, 96% and 118% was accounted for by C content of the macroaggregate fraction at Lakeshore and Gallistel, respectively, under high *Amynthas* pressure vs. control. Of the enriched C in the macroaggregate fraction, about half was attributed to the occluded microaggregate fraction, with the remainder of the C increase presumably in the occluded silt + clay and POM fractions.

**Figure 2.**
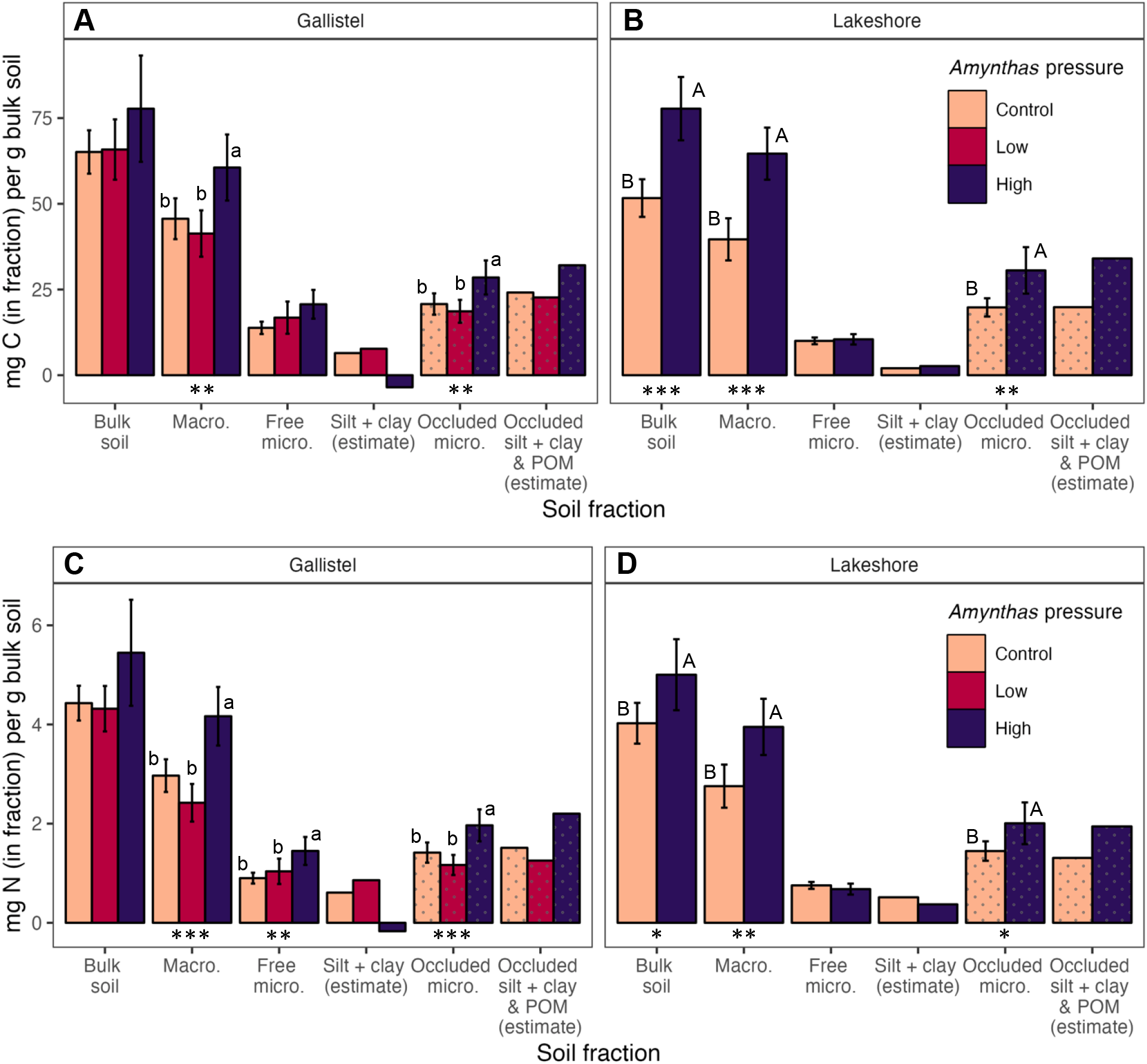
Carbon (**A** & **B**) and nitrogen (**C** & **D)** content in each soil fraction, on a per unit bulk soil basis at Gallistel (**A** & **C**) and Lakeshore (**B** & **D**). Bulk soil = whole soil; Macro. = macroaggregate fraction, 250–2000 µm; Free micro. = microaggregate fraction from bulk soil, 53–250 µm; Silt + clay (estimate) = Carbon content in the < 53 µm fraction, estimated as Bulk soil – (Macro + Free micro); Occluded micro. = microaggregate fraction occluded within macroaggregate fraction, 53–250 µm; Occluded silt + clay & POM (estimate) = Carbon content in the < 53 µm fraction occluded within the macroaggregate fraction, estimated as Macro – Occluded micro. Error bars represent ± 1.96 SE. Asterisks indicate significant treatment differences within soil fraction: *** = *p* < 0.001, ** = *p* < 0.01, * = *p* < 0.05. Bars with different letters (within the same site and fraction) are significantly different (*p* < 0.05). The estimated silt + clay carbon contents do not have associated error bars or statistics. Speckled bars represent occluded fractions.

Contributing to the increased C content of the macroaggregate fraction at Lakeshore, high *Amynthas* pressure significantly increased both the proportion of soil in the macroaggregate fraction compared to the control (Fig. S2B; *p* < 0.01), as well as the C concentration in both macroaggregate and occluded microaggregate fractions (Fig. S3; *p* < 0.01); these metrics also increased under high *Amynthas* pressure at Gallistel, but were not significant. Overall, the occluded microaggregate fraction had a higher C concentration than the free microaggregate fraction, particularly in the high *Amynthas* pressure treatment (*p* < 0.05, Table S2).

N content trends were very similar to those of C (Fig. 2C and D; Fig. S4); N content generally increased under high *Amynthas* pressure, and was attributable to the macroaggregate fraction.

The soil C:N ratio demonstrated significant effects of *Amynthas* pressure (*p* < 0.001) and soil fraction (*p* < 0.05) at both sites (Table S2 and Fig. S5). At Gallistel, C:N ratio was generally widest under low *Amynthas* pressure, followed by the control, then high *Amynthas* pressure, and greater in the macroaggregate fraction compared to the bulk soil. At Lakeshore, the C:N ratio was wider in the high *Amynthas* treatment compared to the control (*p* < 0.001).

High *Amynthas* pressure increased 6h lab-estimated soil respiration on a per unit soil basis and on a per unit soil C basis by 59% and 34%, respectively, at Gallistel (*p* < 0.05; Fig. S6). At Lakeshore, however, high *Amynthas* pressure decreased soil respiration on a per unit soil basis and on a per unit soil C basis by 40% and 62%, respectively (*p* < 0.05; Fig. S6).

### 3.2 *Amynthas* pressure affected bacterial community composition and richness

There was a significant effect of *Amynthas* pressure treatment on bacterial community composition at both sites (R^2^ = 0.18, *p* < 0.001 for Gallistel and R^2^ = 0.16, *p* < 0.001 for Lakeshore, PERMANOVA; Fig. 3). Pairwise testing demonstrated a significant difference between the control and high *Amynthas* treatments at Gallistel and Lakeshore (*p* < 0.001), and between the low and high *Amynthas* treatments at Gallistel (*p* < 0.001). The homogeneity of variance test (BETADISPER) was also significant for *Amynthas* pressure treatment at both sites (*p* < 0.01), which indicates that differences in sample dispersion may have affected results from PERMANOVA. There was also a significant effect of soil pH on bacterial composition, particularly at Gallistel (R^2^ = 0.23, *p* < 0.001 for Gallistel and R^2^ = 0.08, *p* < 0.05 for Lakeshore, PERMANOVA; Fig. S7), which conceivably influenced the sample community composition dispersion. There was no significant effect of soil fraction and no interaction effect of *Amynthas* treatment × soil fraction on community composition at either site. The worm cast samples, which clearly differentiate from the soil samples (Fig. 3; R^2^ = 0.08, *p* < 0.001 for Gallistel and R^2^ = 0.05, *p* < 0.001 for Lakeshore, PERMANOVA), were not included in the above PERMANOVA analyses of the soil communities in order to target compositional differences in the soil.

**Figure 3.**
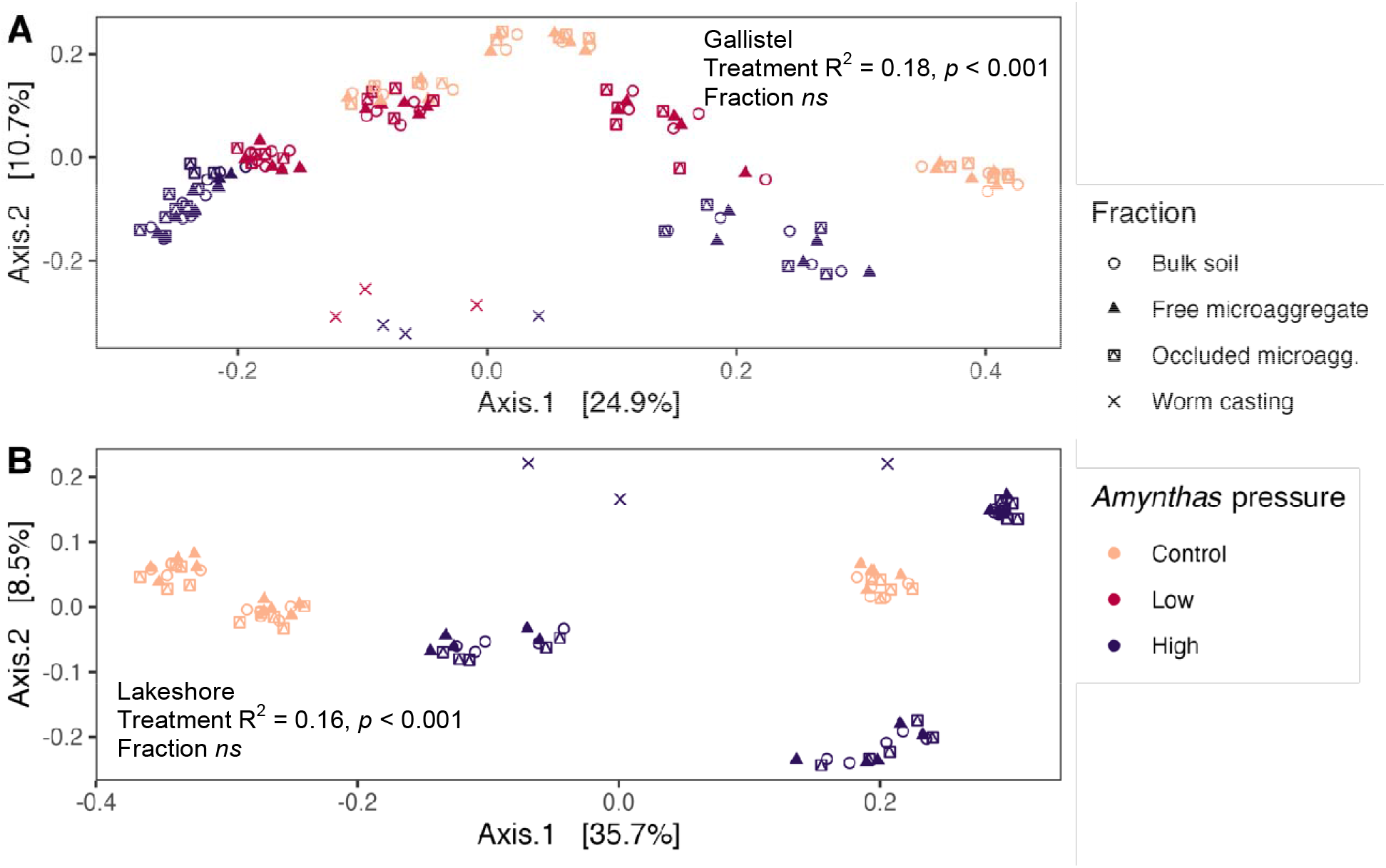
Principal coordinates analysis of Bray-Curtis dissimilarities of Hellinger-transformed community relative abundances, by *Amynthas* pressure treatment for Gallistel (**A**) and Lakeshore (**B**) woodland sites in Madison, WI. Each point represents the community of one sample-fraction. Soil fractions are as follows: Bulk soil = whole soil; Free microaggregate = microaggregate fraction from bulk soil, 53–250 µm; Occluded microagg. = microaggregate fraction occluded within macroaggregate fraction, 53–250 µm; Worm casting = composite casting sample from worms collected in each plot area. Displayed statistics are from PERMANOVA analysis, which excluded worm casting samples.

At Lakeshore, *Amynthas* pressure decreased dispersion of community composition relative to the control (i.e., decreased beta diversity within *Amynthas*-affected soils) by 9% and by 6% at the between-plot and the within-plot scales, respectively, as quantified by mean distance to spatial median (Fig. 4D and E; *p* < 0.01, PERMDISP). The opposite was found at Gallistel, where *Amynthas* pressure increased dispersion by 10-14% at the within-plot scale (Fig. 4B; *p* < 0.001, PERMDISP), with no significant difference at the between-plot scale. The dispersion of the free microaggregate and occluded microaggregate communities within each soil core also increased with *Amynthas* pressure at Gallistel (*p* < 0.001; Fig. 4C) by 12-18%; there was no within-soil core difference in dispersion at Lakeshore (Fig. 4F).

**Figure 4.**
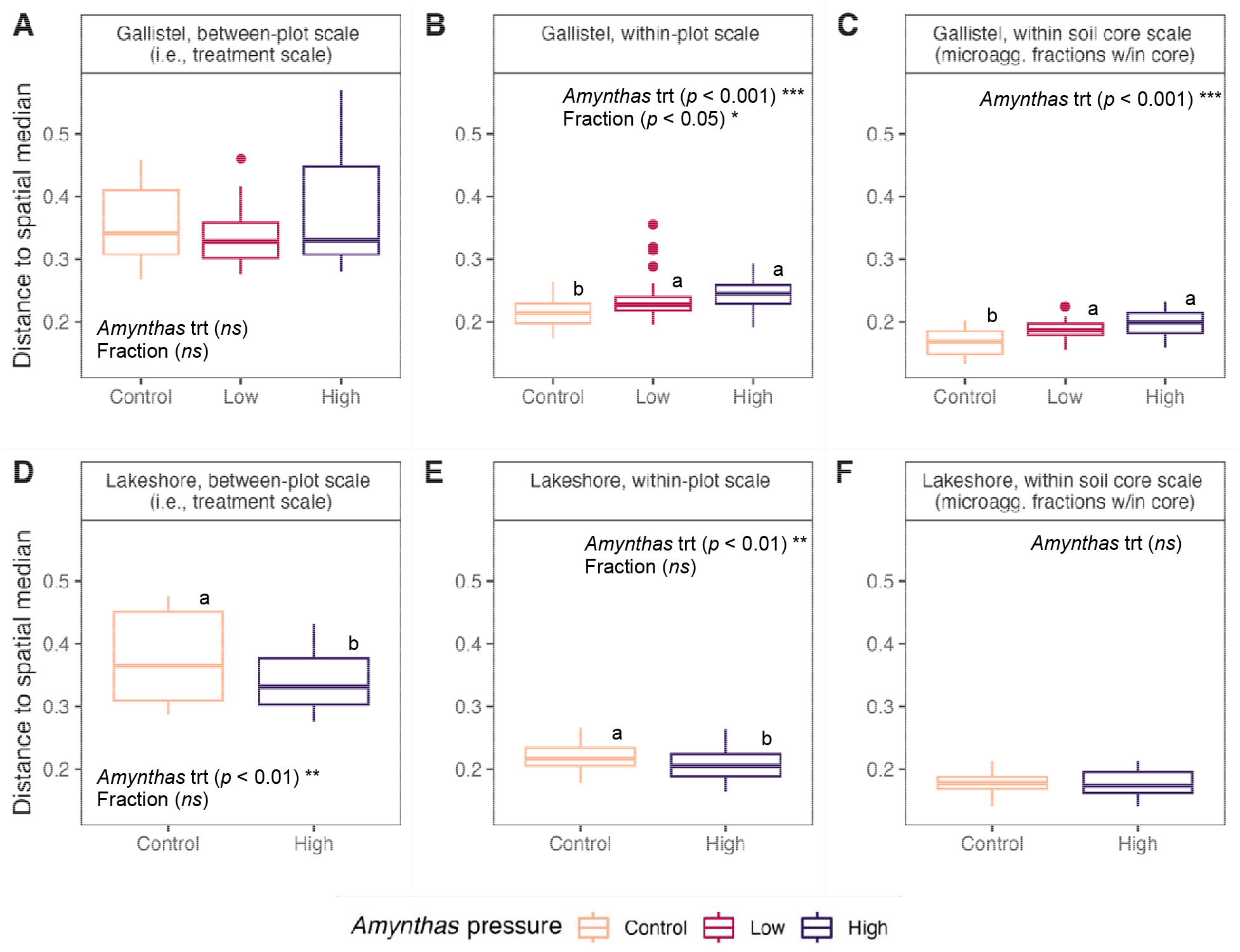
Dispersion of community compositional data as Bray-Curtis dissimilarities, by *Amynthas* pressure treatment, represented as distance to spatial median at the between-plot scale (i.e., treatment scale; **A** and **D**); within-plot scale (**B** and **E**); and soil core scale (free vs. occluded microaggregate fraction samples within a soil core; **C** and **F**) at Gallistel (**A**, **B**, and **C**) and Lakeshore (**D**, **E**, and **F**) woodland sites in Madison, WI. Different letters within the same subfigure denote statically significant treatment differences.

Bacterial OTU richness was significantly affected by *Amynthas* pressure at both sites (*p* < 0.01, Fig. S8). Richness estimates for the control and high *Amynthas* treatments were 8-9% lower compared to the low *Amynthas* pressure treatment at Gallistel (*p* < 0.01, Tukey’s HSD), and richness of the high *Amynthas* treatment was 6% lower than that of the control at Lakeshore. Faith’s PD was also affected by *Amynthas* pressure at both sites (*p* < 0.001, Fig. S9). Faith’s PD for the high *Amynthas* treatment was 5-8% lower than the control or low *Amynthas* treatments at both sites (*p* < 0.05, Tukey’s HSD). There was no significant effect of soil fraction and no interaction effect of *Amynthas* treatment × soil fraction on bacterial richness or Faith’s PD at either site.

### 3.3 Community assembly processes under *Amynthas* pressure

Using the full community comparison approach (Stegen et al., 2015, 2013, 2012), and constructing null models from site-wide data, homogeneous selection dominated community assembly, with > 95% of pairwise comparisons at Gallistel, and 100% of pairwise comparisons at Lakeshore, both within-plot and between-plot, for each set of samples representing *Amynthas* pressure treatment and fraction combinations. This outcome is consistent with the uniformity of plots given constraints within site and *Amynthas* pressure treatment.

Going forward, and in the Discussion section, we will focus on the results from the OTU binning-based approach to characterizing community assembly processes (Ning et al., 2020). Regardless of *Amynthas* pressure treatment, homogenizing dispersal was the dominant community assembly process, followed by homogeneous selection at the within-plot scale (Fig. 5A and C). However, *Amynthas* pressure affected the relative importance of these processes differently at the two sites. At Gallistel, homogenizing dispersal had a mean relative influence of 52% in the control treatment (across fractions), which decreased significantly in low and high *Amynthas* treatments to about 38% (*p* < 0.001 for all fractions) (Fig. 5A). At Lakeshore, homogenizing dispersal had a 51%–58% relative influence across fractions in the control, which increased to about 61% in the high *Amynthas* treatment (*p* < 0.001 for bulk soil and *p* < 0.05 for free microaggregate fractions only) (Fig. 5C). Homogeneous selection was very consistent across fractions at Gallistel, with 15% relative influence in the control, and a significant increase to 21% relative influence in low and high *Amynthas* treatments (*p* < 0.01 for all fractions). At Lakeshore, homogeneous selection in bulk soil decreased from 19% in the control to 14% in the high *Amynthas* treatment (*p* < 0.01), but increased in the occluded microaggregate fraction from 13% in the control to 17% relative influence in the high *Amynthas* treatment (*p* < 0.01), and there was no significant treatment difference in the free microaggregate fraction, with about 14% relative influence.

**Figure 5.**
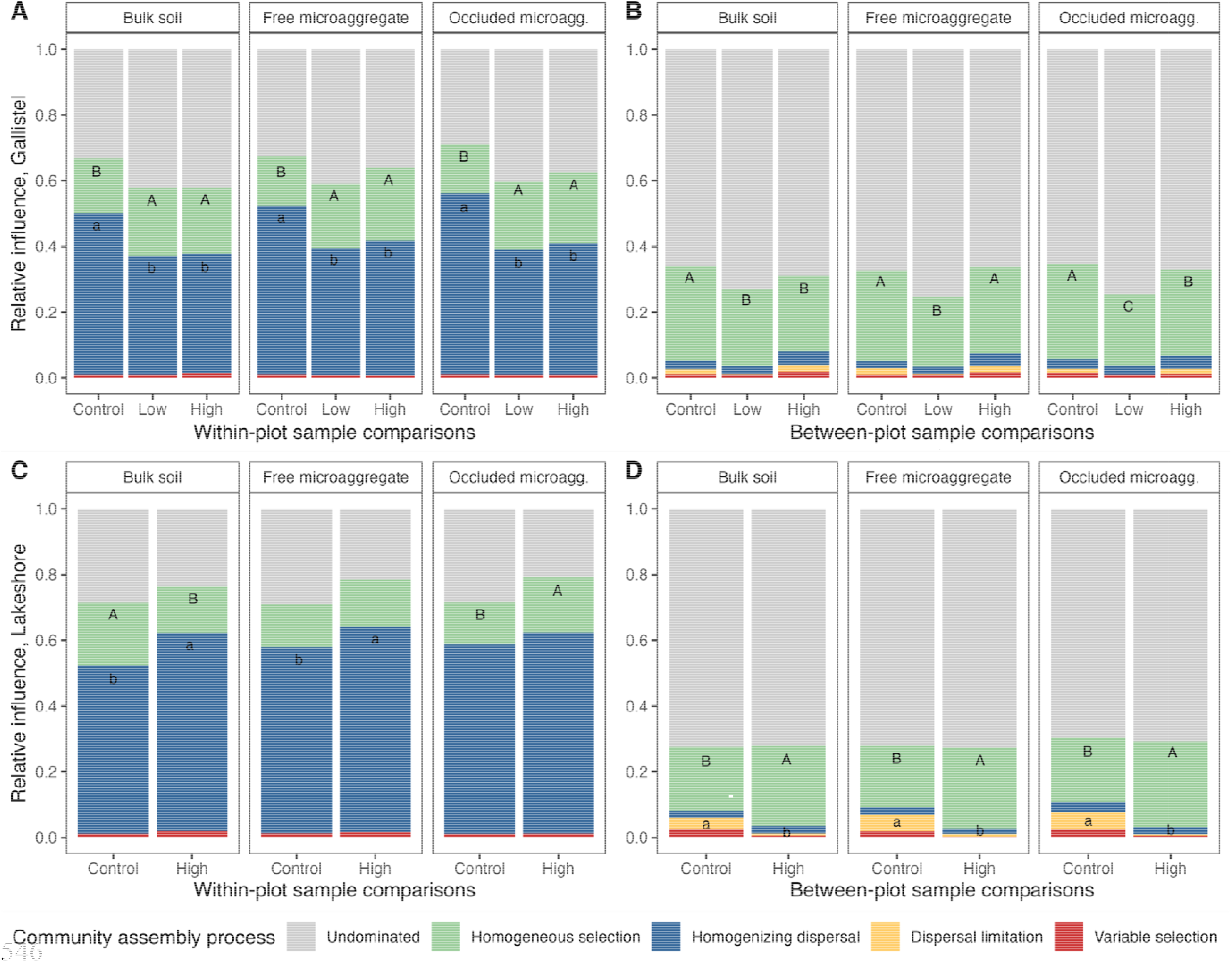
The dominant community assembly processes, by *Amynthas* treatment, within bulk soil, free microaggregate, and occluded microaggregate fractions at Gallistel (**A** and **B**); and Lakeshore (**C** and **D**) woodlands in Madison, WI. Sample comparisons were made within-plot (**A** and **C**) or between-plot (**B** and **D**). Community assembly processes were assigned within phylogenetically related bins of OTUs for pairwise comparisons of samples using a null modeling approach, and weighted by the relative abundance of OTUs in that bin to comprise the full-community (Ning et al., 2020). As detailed in the text, first the influence of selection was determined using the -mean nearest taxon distance, and then the influence of dispersal was determined using the modified Raup-Crick metric based on Bray-Curtis dissimilarity. For the processes with > 5% influence, different letters signify a statistically significant difference in the influence of that process due to *Amynthas* pressure (within site and fraction).

The majority of between-plot comparisons were undominated—over 60% and over 70% at Gallistel and Lakeshore, respectively. Among identified processes, homogeneous selection was dominant, with 29% and 19% relative influence at Gallistel and Lakeshore, respectively, in the control (Fig. 5B and D). This influence generally decreased from 29% in the control to about 24% under *Amynthas* pressure in bulk soil at Gallistel, but increased from 19% to about 25% at Lakeshore across fractions. At Lakeshore, there was also a 5% relative influence of dispersal limitation for between-plot comparisons amongst control samples, which decreased to <1% under high *Amynthas* pressure.

### 3.4 *Amynthas* pressure decreased weighted mean predicted 16S rRNA gene copy number

At both sites, there was a small but statistically significant 2-3% decrease in the weighted mean predicted 16S rRNA gene copy number with *Amynthas* pressure (*p* < 0.001 at Gallistel and *p* < 0.05 at Lakeshore; Fig. 6). Soil fraction was also significant at Gallistel (*p* < 0.05); the weighted mean predicted 16S gene copy number was 2% lower in the occluded microaggregate fraction relative to the bulk soil. There was no significant interaction effect.

**Figure 6.**
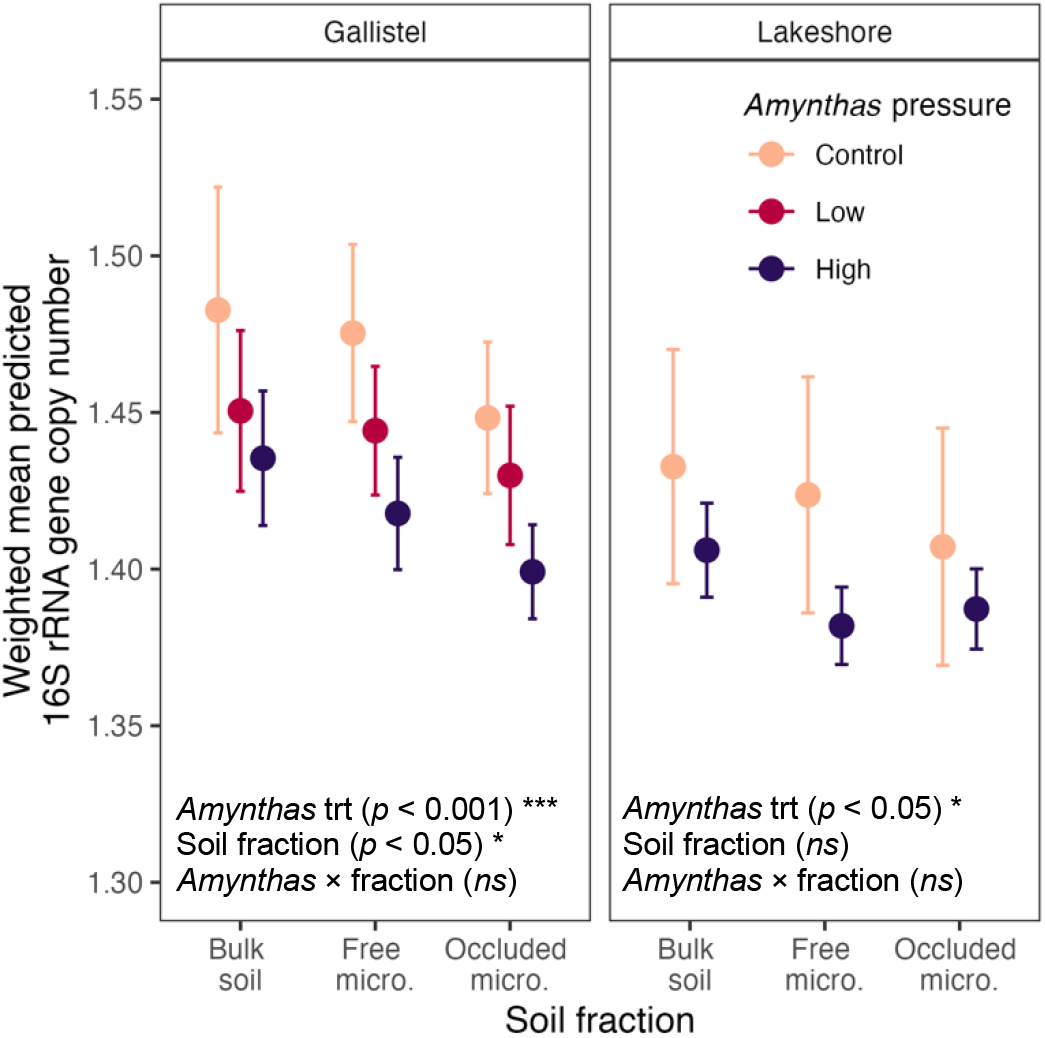
Weighted mean predicted 16S rRNA gene copy number. These data represent taxa for which a gene copy number was available in the rrnDB (Stoddard et al., 2015). Bulk soil = whole soil; Free micro. = microaggregate fraction from bulk soil, 53–250 µm; Occluded micro. = microaggregate fraction occluded within macroaggregate fraction, 53–250 µm. Error bars represent ± 1.96 SE.

### 3.5 Taxonomic differences attributable to *Amynthas* pressure

The most common phyla observed in the soil were *Actinobacteria*, *Proteobacteria, Acidobacteria*, and *Verrucomicrobia*, which together comprised over 70% of relative abundance at both sites. At Gallistel, *Amynthas* pressure resulted in significantly increased relative abundances of *Crenarchaeota* and *Gemmatinomadetes*, significantly decreased relative abundances of *Myxococcota*, and mixed effects across fractions or *Amynthas* pressure levels on *Acidobacteria*, *Bacteroidetes*, and *Verrucomicrobia* (see Fig. S10 for relative abundances and *p* values). At Lakeshore, *Amynthas* pressure resulted in significantly increased relative abundances of *Acidobacteria* and *Methylomirabilota*, and significantly decreased relative abundances of *Gemmatimonadetes* and *Proteobacteria* (see Fig. S11 for relative abundances and *p* values).

We also identified key taxa associated with *Amynthas* pressure, based on differential abundance. Across both sites, we identified a total of 1283 taxa that were enriched in *Amynthas* treatments (relative to control), and 1285 taxa that were depleted in *Amynthas* treatments. We focused on the taxa with the biggest responses (|µ| > 1.0), and only considered taxa with mean relative abundances greater than 0.001 (0.1%), which resulted in 34 and 21 focal taxa enriched at Gallistel and Lakeshore, respectively (Fig. S12). Though some taxa were unique responders within a soil fraction, there were numerous taxa that were enriched across bulk soil, free microaggregate, and occluded microaggregate fractions, including ubiquitous *Actinobacteria* such as *Mycobacterium* and *Gaiella*. At Gallistel, there was some enrichment of anaerobic taxa (e.g., *Anaerolineae*) (Yamada and Sekiguchi, 2020), and at Lakeshore, there were several enriched *Nitrososphaera* taxa (*Crenarchaeota*), which are aerobic ammonia oxidizers (Stieglmeier et al., 2014), perhaps taking advantage of *Amynthas*-driven increases in ammonium concentrations (Qiu and Turner, 2017). There was only one focal taxon at each site that was depleted under *Amynthas* activity, and both of these taxa were affiliated with *Candidatus* Udaeobacter (*Verrucomicrobia*), which are common in soil (Brewer et al., 2016).

We looked for taxa that were enriched in the microaggregate fractions relative to bulk soil using differential abundance, however, we found no positively enriched taxa.

### 3.6 Bacterial composition of *Amynthas* castings

The most abundant phyla in worm castings across both sites were *Proteobacteria*, *Bacteroidetes*, *Actinobacteria*, *Firmicutes*, which together comprised over 80% of the relative abundance of OTUs in worm castings from each plot. The most prevalent cast-associated OTU across plots was of the genus *Aeromonas* (*Gammaproteobacteria*), and comprised a mean 16% of relative abundance in casting samples from Lakeshore and a mean 9% of relative abundance in casting samples from Gallistel. However, this OTU was not found to be relatively enriched in the soil communities under *Amynthas* pressure at either site. Only one abundant worm casting OTU (with > 1% relative abundance in any casting sample) from each site was identified as enriched in soil communities: a *Flavobacterium* (*Bacteroidetes*) at Gallistel, and an *Enterobacteriaceae* (*Gammaproteobacteria*) at Lakeshore, and both of these OTUs were still considered ‘rare’ in the soil communities following enrichment (< 0.1% relative abundance).

Because individual worm casting OTUs were generally not enriched in soil communities, we considered an aggregated group of OTUs that represented the top 75% of the relative abundance of worm casting communities at each site, and found the total relative abundance for these groups in soil communities. We found the worm casting OTUs comprised about 14% and 19% of the relative abundance in the soil communities from control plots at Gallistel and Lakeshore, respectively (i.e., present in the native soil community prior to *Amynthas* invasion), with a significant increase in the relative abundance of these site-specific groups of OTUs in high *Amynthas* pressure plots at Gallistel only, to about 18% relative abundance in the microaggregate fractions (Fig. S13).

## 4. Discussion

We sought to test the effects of soil physical disturbance due to the activity of non-native *Amynthas tokioensis* and *Amynthas agrestis* earthworms on the soil bacterial community, specifically in soil microaggregate fractions. Activity of these *Amynthas* spp. earthworms restructures the soil surface, creating a layer of worm castings. Despite a high level of *Amynthas* activity that presumably constituted thorough soil mixing, the microbial community composition did not consistently reflect the effects of homogenization that have been previously observed under soil mixing-type disturbances.

### 4.1 *Amynthas* pressure increased total C and N, with contrasting effects on soil respiration

Given previous evidence for C and N increases in surface soil at this site (Qiu and Turner, 2017), and elsewhere under *A. agrestis* invasion (Zhang et al., 2013), the enrichment of C and N in the macroaggregate and occluded microaggregate fractions under high *Amynthas* pressure (Fig. 2) supports the idea that earthworm-enhanced aggregate formation is a mechanism for short-term increases in C and N in surface soil. That said, this finding is interpreted with caution because numerous studies suggest that earthworm-accelerated litter decomposition may reduce total C stocks over time (Chang et al., 2017; Fahey et al., 2013; Qiu and Turner, 2017), and previous work at Gallistel found increased C metabolic diversity shortly (∼ 2 years) after invasion using community-level physiological profiling, indicating shifts in patterns of microbial C metabolism (Price-Christenson et al., 2020).

Soil respiration, measured from sieved soil in the lab, demonstrated contrasting responses to *Amynthas* pressure between the two sites (Fig. S6). This may be attributable to differences in the length of time since invasion, which is not definitively known for the high *Amynthas* pressure treatment at either site (see section 2.1, Soil collection, for details on worm survey data). We do know that the low *Amynthas* pressure plots at Gallistel had been under invasion for no more than four years at the time of sampling, and demonstrated an increase in soil respiration rate, particularly on a per unit of soil C basis. This corroborates previously measured increases in respiration following recent invasion at this site (Price-Christenson et al., 2020), indicating a stimulation of microbial activity, likely due to increased resource availability associated with *Amynthas* activity and litter incorporation. As length of time since invasion increases, resources may become depleted or inaccessible to microbes once under aggregate protection, or the microbial community may decrease due to earthworm consumption (Chang et al., 2016a), resulting in decreased respiration, as was measured at Lakeshore and trending at Gallistel under high *Amynthas* pressure (Fig. S6B). Invasion-driven changes in the soil microbial community over time may also affect soil respiration rates (McLean et al., 2006). The small decrease in weighted mean predicted 16S rRNA gene copy number under *Amynthas* pressure (Fig. 6) might suggest a shift towards a community that uses C more efficiently (Roller et al., 2016), with an associated decrease in respiration per unit of soil C. Given increased resource availability under epi-endogeic earthworm activity, and previous work that demonstrates increased potential for fast growth under soil disturbance (West et al., 2023; West and Whitman, 2022; Whitman et al., 2019), we did not expect the decrease in weighted mean predicted 16S rRNA gene copy number under *Amynthas* pressure, however small. Formation of dense, anoxic aggregates via *Amynthas* activity perhaps detracted from potential benefits of fast growth, while favoring facultative or anaerobic community members.

Different sample collection dates may also explain contrasting respiration responses between sites. A burst of microbial activity at Gallistel could have been driven by increased N availability (observed in Qiu and Turner, 2017) from earthworm mortality itself (Chang et al., 2021), whereas Lakeshore was sampled two weeks later, and might instead reflect microsite resource depletion in the lag time since *Amynthas* die-off and cessation of bioturbation activity, as evidenced by decreased respiration. Though we did not confirm the date of *Amynthas* die-off, it occurred between the 18 October, 2021 worm collection date and the 30 October, 2021 Gallistel soil sampling date, when overnight temperatures in Madison approached freezing; Lakeshore was sampled on 13 November, 2021.

The high sand content in the high *Amynthas* treatment at Lakeshore (44%; Table S1) may have effectively diluted the respiration rate relative to soil from the control plots (22% sand), thus resulting in an overall decrease in measured respiration under *Amynthas* pressure (per unit of soil; Fig. S6A), despite a large increase in soil carbon content (Fig. 2B, Fig. S3). The two treatment sampling areas represent different soils, but the high sand content in the high *Amynthas* area was unexpected based on soil maps. Differences in soil texture is a common feature of field sampling in natural settings, and the > 800 m distance between treatment sampling areas was partially necessary due to the rapid spread of the *Amynthas* invasion. Though a higher sand content may be indicative of lower soil moisture content and thus reduced respiration (Moyano et al., 2012), our data do not support this mechanism, perhaps due to a tradeoff for increased soil carbon content under high *Amynthas* pressure (Fig. S14B). Yet another explanation might be a high proportion of anaerobic microsites in the Lakeshore high *Amynthas* soil—anecdotally, the layer of worm castings was very thick at this site, perhaps comprising the entire 3 cm depth sampled, and it has been demonstrated that highly compacted worm castings contain small pores and anoxic microhabitats (Blanchart et al., 1993; Jouquet et al., 2008). Further, soil with higher sand content and particulate organic matter inputs has been shown to harbor more anoxic microsites compared to soil with higher clay content, due to the increased potential for rapid, oxygen-depleting microbial activity given lower propensity for protective organo-mineral associations associated with the higher sand content (Lacroix et al., 2022).

Overall, we found a strong relationship between C concentration and soil moisture at each site (Fig. S14A) and within each treatment, with the strength and significance of that relationship increasing under *Amynthas* pressure (Fig. S14B). This suggests that increased C content – indicative of SOM enrichment under *Amynthas* pressure – increased soil moisture. Increased soil moisture alone is a known factor to increase soil microbial activity and thus respiration (Davidson et al., 1998), but when we control for carbon content, we see no relationship between soil moisture and respiration at Gallistel (Fig. S15). From this, we conclude that *Amynthas* activity increased SOC, which increased soil microbial respiration, regardless of the effect on soil moisture.

Despite visibly evident *Amynthas*-driven creation of soil aggregates via casting activity, there was no clear trend in the proportion of soil in aggregate fractions (Fig. S1). Though previous work has found *Amynthas* activity (Greiner et al., 2012; Snyder et al., 2011) and earthworm activity in general (Mummey et al., 2006) to increase aggregation, we only observed this at Lakeshore, with an 8% increase. The lack of aggregation response may be attributable to the high baseline level of aggregation at these forested sites, with 70–75% of soil (dry mass basis) in water-stable macroaggregate or free microaggregate fractions in the control treatments (Fig. S1A & S1B). Further, our choice to quantify macroaggregates sized 250 µm–2000 µm potentially missed changes to large macroaggregates > 2000 µm, which might increase under the activity of *Amynthas* or other earthworms (Bossuyt et al., 2006; Knowles et al., 2016; Zhang et al., 2013).

### 4.2 Homogenizing dispersal influences bacterial community assembly at the plot scale

Community assembly findings at Lakeshore generally support hypothesis H1 in that *Amynthas* pressure homogenized soil bacterial communities at the within-plot scale via homogenizing dispersal (Figs. 4E and 5C), and at the between-plot scale via homogeneous selection (Figs. 4D and 5D), thus apparently promoting soil mixing and community coalescence. A moderate influence of dispersal limitation was also noted for between-plot comparisons in the control (also supporting H1). These findings reflect the patterns found in a complementary experiment in which we explored the effects of long-term tillage as a modality of soil mixing (West et al., 2023).

On the other hand, findings at Gallistel do not support our hypothesis, and run contrary to the idea that bioturbation via *Amynthas* pressure acts as a homogenizing force. Here, community compositional sample dispersion at the within-plot scale increases under *Amynthas* pressure (Fig. 4B), perhaps attributable to the decreased influence of homogenizing dispersal (Fig. 5A). There was a slight but significant within-plot increase in homogeneous selection under *Amynthas* pressure, though this trend reversed at the between-plot scale. These inconsistencies, particularly the counter-intuitive decrease in homogenizing dispersal at the within-plot scale, may reflect the variable proportions of soil and worm casts in the *Amynthas*-affected samples (*up to* 3 cm of worm casts plus remainder of depth comprised of soil) vs. control samples (3 cm of soil). A variable proportion of worm castings within a treatment may highlight the inherent differences in composition and community assembly in soil vs. worm castings, as opposed to true treatment differences. Future work might attempt to manually quantify the proportion of worm-aggregated soil in a given sampling depth (Lavelle et al., 2020), or consider taking a mass-equivalent sampling approach rather than a depth- or volume-equivalent approach. The small-scale heterogeneity under *Amynthas* pressure is also consistent with a high level of isolation amongst communities caused by discontinuous pore networks in soil extensively affected by worm casting activity, attributable to a surface barrier on the earthworm-generated aggregates, which forms as water is drawn out of the casting by the worm’s intestine prior to excretion, pulling fine particles to the casting surface to form a cortex (Blanchart et al., 1993).

The overall inconsistencies in community assembly process response to *Amynthas* pressure corroborate other research that found the worm gut microbiota to lack a clear and direct influence on the soil microbial community (McLean et al., 2006; Price-Christenson et al., 2020). If the gut microbiota of *A. tokioensis* and *A. agrestis* earthworms had a strong, direct impact on the microbial communities of invaded soil, we would expect that gut community members would drive increases in both homogenizing dispersal and selection. Instead, the *Amynthas* activity has an indirect effect on the microbial communities, via changes to soil structure and resource availability. Our worm casting data reflect this, since we did not see major markers of worm casting OTU enrichment in *Amynthas*-affected soil communities, and at Gallistel, where we did observe the dominant worm casting community increase in the soil (Fig. S13), there was no concomitant increase in homogenizing community assembly processes.

### 4.3 Evidence for fluidity between the free and occluded microaggregate fractions

We hypothesized that free and occluded microaggregate fractions would support more similar bacterial communities under *A. tokioensis* and *A. agrestis* co-invasion activity compared to the control, due to homogenizing bioturbation and the rapid formation of aggregates via casting formation. However, community composition, OTU richness, and Faith’s PD were unaffected by soil fraction or an interaction effect at either site (Figs. 3, S8, S9). Community assembly processes also trended similarly regardless of free vs. occluded microaggregate fraction, and largely reflected community assembly of the bulk soil communities. At the finest scale, there was even some indication of increased heterogeneity between the free and occluded microaggregate communities under *Amynthas* activity, as indicated by increased dispersion between the free and occluded microaggregate communities within singular soil cores at Gallistel (Fig. 4C). Though, we hesitate to over-interpret the within-core dispersion finding without fraction-wide compositional changes.

*Amynthas agrestis*, and earthworm activity in general, has been associated with increases in the large macroaggregate pool (> 2000 µm) (Bossuyt et al., 2006; Knowles et al., 2016; Zhang et al., 2013), which we did not specifically measure since we sieved soil to < 2000 µm prior to fractionation. Due to potential disruption of large macroaggregates during this initial sieving, some portion of occluded microaggregates may have been isolated in the free microaggregate pool, thus blurring potential differences between the occluded vs. free microaggregate pools. Though this likely occurred to some extent, the significantly higher concentration of soil C in the occluded compared to the free microaggregate fraction (Table S2), particularly under high *Amynthas* pressure, first indicates distinction between these fractions, and, moreover, suggests some environmental protection of C under occlusion, potentially attributable to anoxic conditions created via worm casting activity (Lacroix et al., 2022), despite apparent similarities in community composition between the microaggregate fractions.

### 4.4 Factors that may have diminished the measurable impacts of *Amynthas* spp. activity

*Amynthas* spp. were not the first earthworms to invade these woodlands, and the effects of bioturbation and casting activity of the non-native European earthworm precursors may be apparent, particularly in the control plots. Otherwise, spatial variability of edaphic characteristics and vegetation are challenging factors to reconcile in long-term, non-manipulative field work. These factors are further complicated under co-invasion, because the different species will display variable feeding habitats and casting characteristics depending on environment and resource availability, as well as cross-species interactions (Chang et al., 2018; Richardson et al., 2022). Though the precise length of time since invasion in the high *Amynthas* pressure plots is unknown, we can reasonably speculate that the major effects of the invasion began around 2013, when these earthworms were first noted in the Arboretum. The severe drought in southern Wisconsin in 2012 likely would have profoundly affected any potential existing population, causing a retraction in the extent of invasion, or reduction in abundance or fecundity (e.g., cocoon production) due to drought-related energy expenditures (Snyder et al., 2013, 2011).

## 5. Conclusions

We assessed bacterial community composition of free and occluded microaggregate fractions in soil under co-invasion by *Amynthas tokioensis* and *Amynthas agrestis*. Instead of focusing on the characteristics and microbial community impacts due to physical soil disturbance under earthworm bioturbation, as was the original motivation for this work, we instead identified unique impacts of the co-invasion by *A. tokioensis* and *A. agrestis* specifically attributable to these earthworms’ ecology. Though the soil aggregates created through *Amynthas* spp. casting behavior may be operationally defined by standard aggregate fraction isolation techniques (e.g., wet sieving), mixed evidence for homogenizing forces indicates that these aggregates are not well-integrated into the soil matrix, and instead promote heterogeneous microbial community compositions due to their dense, non-porous structure that fosters anoxic microhabitats. It would be valuable to visualize the architecture of these earthworm-derived aggregates, and quantify pore network characteristics, in order to better describe the microhabitats within. Field assessments of soil respiration using isotope-labelled litter would also be valuable in order to better predict carbon transformations and evaluate tradeoffs between litter and SOM consumption vs. the potential protections conferred through worm casting formation.

## Supplementary Information

Supplementary Information can be found online.

## Acknowledgements

The authors would like to thank Alexa Hanson, Kallysa Taylor, Emma Johnson, Isabelle Bartholomew, and Eren Wolf for their indispensable help as undergraduate assistants; Dana Johnson and the other members of the Whitman lab for their thoughtful input; Daliang Ning for guidance with iCAMP analysis; Harry Read and Anna Cates for their perspectives on soil fractionation and use of the microaggregate isolator; and Katie Laushman for initial work establishing plots at the UW-Arboretum.

The authors also acknowledge the UW Biotechnology Center DNA Sequencing Facility (Research Resource Identifier—RRID:SCR_017759) for providing library sequencing facilities and services. Part of this research was performed using the computational resources and assistance of the UW-Madison Center for High Throughput Computing (CHTC; https://doi.org/10.21231/GNT1-HW21), with the help of Christina Koch. The CHTC is an active member of the OSG Consortium, which is supported by the National Science Foundation (NSF) and the U.S. Department of Energy’s Office of Science. This work was financially supported by the O.N. Allen Professorship (UW-Madison CALS), the Louis and Elsa Thomsen Wisconsin Distinguished Graduate Fellowship (UW-Madison CALS), a NSF EAGER grant (award #2024230), and the Wisconsin Department of Natural Resources.

## Author contributions

JW and TW conceived of the project. BH established and monitored plots in the Arboretum and Lakeshore Nature Preserve since 2015 and 2017, respectively, thus providing historical data to leverage toward plot selection. JW collected soil samples, conducted lab work, analyzed the data, and drafted the manuscript. All authors reviewed and edited the manuscript.

## Abbreviations

(SOM): soil organic matter
(SOC): soil organic carbon
(POM): particulate organic matter

