## Supplementary_Information for "Soil microaggregate bacterial communities following *Amynthas tokioensis* and *Amynthas agrestis* earthworm co-invasion"

### **Method: PCR amplification, and 16S rRNA library prep**

All DNA was stored at or below −20 °C throughout the experiment. Each round of soil extraction included one extraction blank, which was carried through the following pipeline alongside soil samples. 16S rRNA genes of extracted DNA were amplified in triplicate using PCR. Variable region V4 of the 16S rRNA gene was targeted using forward primer 515f and reverse primer 806r (Walters et al., 2016). Primers also contained barcodes and Illumina sequencing adapters (Kozich et al., 2013). The following reagents comprised each 25 μL PCR reaction: 12.5 μL Q5 Hot Start High-Fidelity 2X Master mix (Catalog No. M0494, New England BioLabs, Ipswich, MA), 1.25 μL 515f forward primer (10 mM), 1.25 μL 806r reverse primer (10 mM), 1 μL DNA extract, 1.25 μL Bovine Serum Albumin (20 mg/mL; Catalog No. 97064-342, VWR International, Radnor, PA, and 7.75 μL PCR-grade water. The plate was sealed and briefly centrifuged prior to 30 PCR cycles on an Eppendorf Mastercycler nexus gradient thermal cycler (Hamburg, Germany) using the following parameters: 98 °C for 2 min + 30 × (98 °C for 10 seconds + 58 °C for 15 seconds + 72 °C for 10 seconds) + 72 °C for 2 min and 4 °C hold. Successful amplification was verified via gel electrophoresis using 1% TAE agarose gel and Invitrogen SYBR Safe DNA Gel Stain (Catalog No. S33102, ThermoFisher Scientific, Carlsbad, CA). Each well was loaded with 5 μL PCR product mixed with 1 μL Purple 6X Gel Loading Dye (Catalog No. B7025S, New England BioLabs, Ipswich, MA). Successful amplification of target base pair length was confirmed using a 2-Log DNA Ladder (0.1–10.0 kilobases; Catalog No. N3200, New England BioLabs, Ipswich, MA) in each gel. Gel ran for ninety minutes at 115V. Gels were photographed and checked visually for amplification.

The SequalPrep Normalization Plate Kit (Catalog No. A1051001, ThermoFisher Scientific, Carlsbad, CA) was used to normalize amplicon yields across samples using a limited binding capacity solid phase. Normalization was performed following kit instructions using 25 μL of the pooled triplicate PCR product, and yielded 20 μL of eluted DNA per sample, which was pooled following elution. The combined DNA library was concentrated using a SpeedVac Vacuum Concentrator System (ThermoFisher Scientific, Waltham, MA) prior to further DNA purification using Wizard SV Gel and PCR Clean-Up System (Catalog No. A9281, Promega Corporation, Madison, WI). Kit instructions were followed except the nuclease-free water was divided into 30 μL and 20 μL increments with the incubation and centrifuge steps after both additions.

### Supplementary Table

**Table S1**. Soil properties based on representative subsamples for each treatment at each site, 0-3 cm. OM = organic matter; CEC = cation exchange capacity. Soil texture (percent sand, silt, and clay) was determined via hydrometer method (Bouyoucos, 1962); organic matter was determined via loss on ignition (Schulte and Hopkins, 1996); pH was determined for 1:1 water (Richards, 1954); plant-available P and K were determined by Bray-1 method (Bray and Kurtz, 1945); and plant-available Ca and Mg were determined by ammonium acetate method (Thomas, 1982).

| **Site** | **Treatment** | **Sand** | **Silt** | **Clay** | **OM** | **Texture** | **pH** | **P** | **K** | **Ca** | **Mg** | **CEC** |
|---|---|---|---|---|---|---|---|---|---|---|---|---|
| | | ----------------- *%* ----------------- | | | | | | ---------------- *mg kg$^{-1}$* ---------------- | | | | |
| **Gallistel** | **Control** | 24 | 58 | 18 | 9.6 | Silt loam | 5.9 | 36 | 109 | 1904 | 380 | 18 |
| | **Low *Amynthas*** | 26 | 58 | 16 | 8.1 | Silt loam | 6.1 | 46 | 87 | 2445 | 337 | 19 |
| | **High *Amynthas*** | 26 | 58 | 16 | 9.7 | Silt loam | 6.2 | 22 | 96 | 2474 | 304 | 20 |
| **Lakeshore** | **Control** | 22 | 58 | 20 | 7.5 | Silt loam | 6.4 | 83 | 150 | 2331 | 481 | 20 |
| | **High *Amynthas*** | 44 | 40 | 16 | 8.6 | Loam | 6.9 | 77 | 131 | 2372 | 548 | 19 |

**Table S2**. Soil carbon and nitrogen concentrations and C:N ratio in soil fractions, by *Amynthas* pressure treatment, presented as mean and (standard error) with ANOVA *p* values. Means followed by the same letter (within site) are not statistically different. Bulk soil = whole soil; Macroaggregate = 250–2000 µm; Free microaggregate = microaggregate fraction from bulk soil, 53–250 µm; Occluded microagg. = microaggregate fraction occluded within macroaggregate fraction, 53–250 µm.

| | | Gallistel | | | Lakeshore | |
|---|---|---|---|---|---|---|
| | *Soil fraction* | Control | Low *Amynthas* | High *Amynthas* | Control | High *Amynthas* |
| Soil total carbon concentration (%C) | Bulk soil | 6.5 (0.32) d | 6.6 (0.45) d | 7.8 (0.79) cd | 5.2 (0.28) E | 7.8 (0.47) BCD |
| | Macroaggregate | 8.7 (0.44) cd | 9.8 (0.68) bcd | 11.1 (0.89) abc | 6.1 (0.37) DE | 8.8 (0.41) BC |
| | Free micro. | 8.8 (0.70) cd | 10.5 (1.1) abc | 8.7 (0.94) cd | 7.0 (0.45) CDE | 7.2 (0.65) BCD |
| | Occluded micro. | 11.0 (0.52) abc | 13.6 (0.81) a | 13.2 (0.89) ab | 9.1 (0.41) B | 11.4 (0.54) A |
| | *Source of variation* | | *p* value | | *p* value | |
| | *Amynthas* treatment | | < 0.01 ** | | < 0.001 *** | |
| | Soil fraction | | < 0.001 *** | | < 0.001 *** | |
| | *Amynthas* × fraction | | 0.2260 | | < 0.05 * | |
| Soil total nitrogen concentration (%N) | Bulk soil | 0.44 (0.02) e | 0.43 (0.02) e | 0.54 (0.05) de | 0.40 (0.02) C | 0.50 (0.04) C |
| | Macroaggregate | 0.56 (0.03) de | 0.57 (0.03) cde | 0.76 (0.05) abc | 0.42 (0.02) C | 0.54 (0.03) BC |
| | Free micro. | 0.57 (0.04) cde | 0.66 (0.06) bcd | 0.61 (0.06) cde | 0.53 (0.04) BC | 0.48 (0.05) C |
| | Occluded micro. | 0.75 (0.03) abcd | 0.84 (0.04) ab | 0.91 (0.05) a | 0.66 (0.03) AB | 0.75 (0.03) A |
| | *Source of variation* | | *p* value | | *p* value | |
| | *Amynthas* treatment | | < 0.001 *** | | < 0.05 * | |
| | Soil fraction | | < 0.001 *** | | < 0.001 *** | |
| | *Amynthas* × fraction | | 0.1702 | | 0.0693 | |
| C:N ratio | Bulk soil | 14.6 (0.20) d | 15.1 (0.36) bcd | 14.3 (0.19) d | 12.9 (0.28) E | 16.1 (0.81) AB |
| | Macroaggregate | 12.2 (0.25) bcd | 17.0 (0.34) a | 14.5 (0.40) d | 14.4 (0.26) BCDE | 16.8 (0.63) A |
| | Free micro. | 13.6 (0.17) bcd | 15.9 (0.43) abc | 14.2 (0.14) d | 13.3 (0.21) DE | 15.8 (0.70) ABC |
| | Occluded micro. | 13.3 (0.11) cd | 16.0 (0.28) ab | 14.4 (0.20) d | 13.7 (0.17) CDE | 15.3 (0.32) ABCD |
| | *Source of variation* | | *p* value | | *p* value | |
| | *Amynthas* treatment | | < 0.001 *** | | < 0.001 *** | |
| | Soil fraction | | < 0.01 ** | | < 0.0451 * | |
| | *Amynthas* × fraction | | 0.0471 * | | 0.4081 | |

**Table S3**. Taxa enriched in low *Amynthas* and high *Amynthas* pressure treatments, by fraction. Includes sequences and statistics from differential abundance output. *See accompanying file SI_Table_S3_enriched_taxa_by_Amynthas_pressure_trt_and_soil_fraction.csv*

### Supplementary Figures

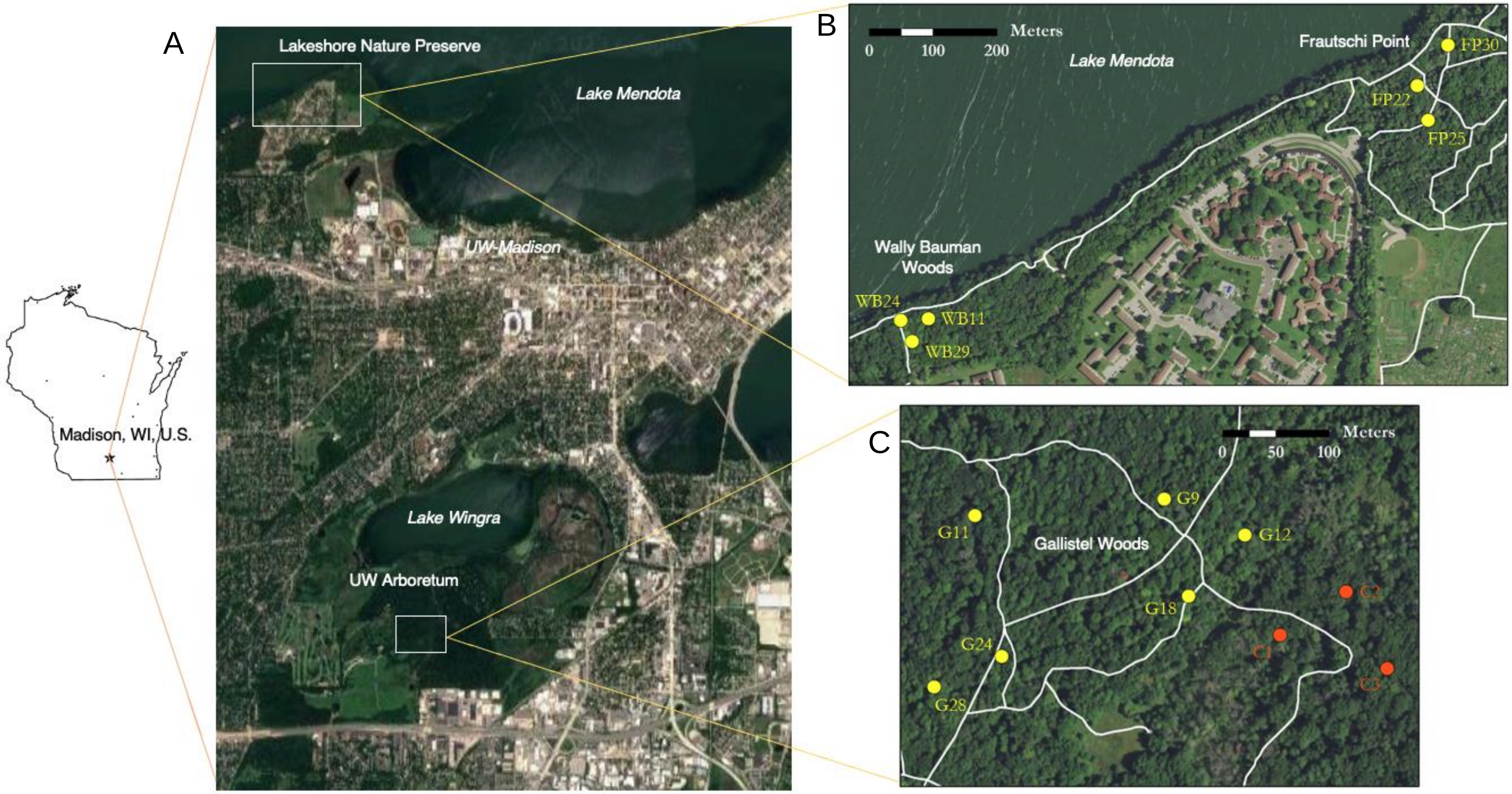

**Figure S1**. Maps of plot locations in Madison, WI, U.S. (**A**). The Lakeshore Nature Preserve inset (**B**) details plots in Wally Bauman Woods (high *Amynthas* pressure plots) and Frautschi Point (control). The UW-Arboretum inset (**C**) details plots in Gallistel Woods, with the control plots marked in orange. Hiking trails are in white. Figure **A** was modified from Google maps (4 April, 2023) and **B** & **C** were created by Danielle Tanzer using ArcGIS. Katie Laushman and Brad Herrick established the plots at the UW Arboretum, and Brad Herrick established the plots at the Lakeshore Nature Preserve.

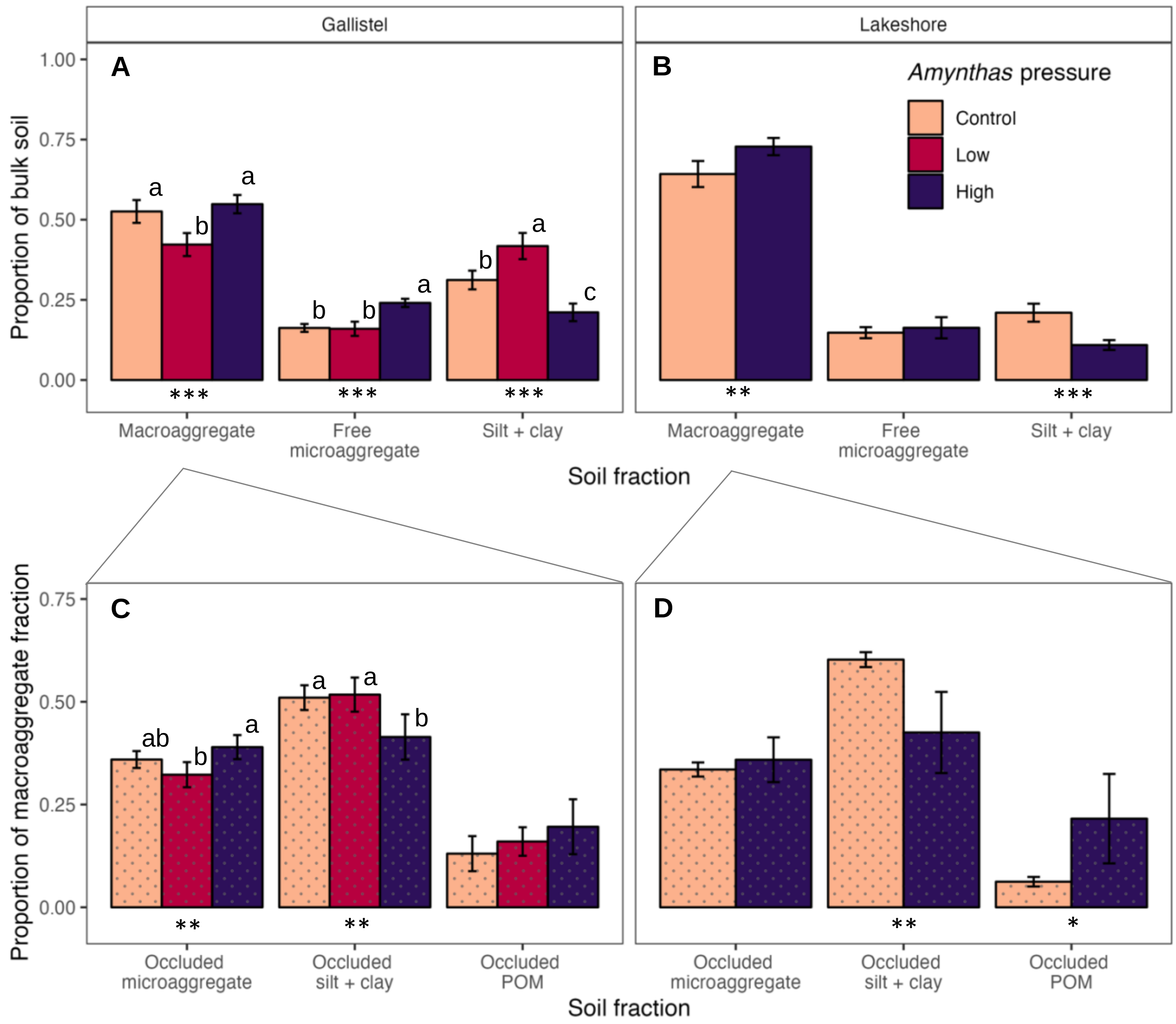

**Figure S2**. Distribution of bulk soil in various fractions at Gallistel (**A**) and Lakeshore (**B**) woods in Madison, WI, on a dry soil basis. Lower panels show distribution of macroaggregate soil in the occluded fractions (**C** and **D**). Macroaggregate = macroaggregate fraction, 250–2000 µm; Free microaggregate = microaggregate fraction from bulk soil, 53–250 µm; Silt + clay = silt and clay-sized fraction from bulk soil, <53 µm; Occluded microaggregate = microaggregate fraction occluded within macroaggregate fraction, 53–250 µm; Occluded silt + clay = silt and clay-sized fraction occluded within the macroaggregate fraction, <53 µm; Occluded POM = particulate organic matter and sand occluded within the macroaggregate fraction, 250–2000 µm. Error bars represent ± 1.96 SE. Asterisks indicate significant *Amynthas* pressure treatment differences within soil fraction: *** = $p < 0.001$, ** = $p < 0.01$, * = $p < 0.05$. Letters signify significantly different treatment within fraction for Gallistel site. Speckled bars represent occluded fractions.

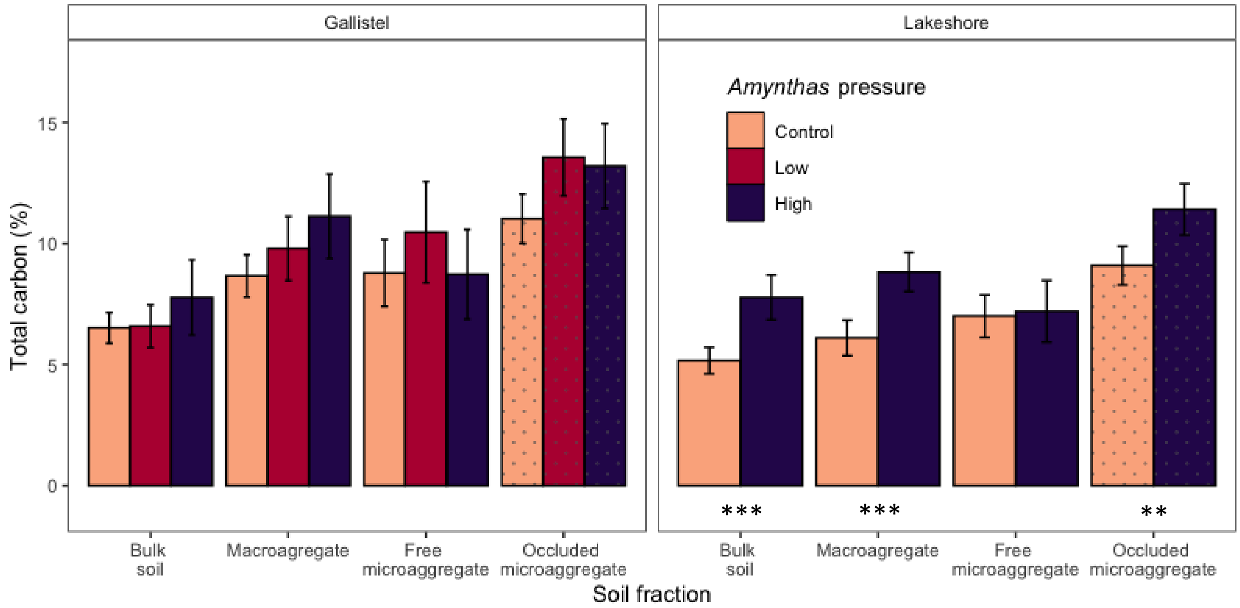

**Figure S3**. Carbon concentration (total C, %), by soil fraction and *Amynthas* pressure treatment. Soil fractions are as follows: Bulk soil = sieved field-moist soil < 2000 µm; Macroaggregate = macroaggregate fraction, 250–2000 µm; Free microaggregate = microaggregate fraction from bulk soil, 53–250 µm; Occluded microaggregate = microaggregate fraction occluded within macroaggregate fraction, 53–250 µm. Error bars represent ± 1.96 SE. Asterisks indicate significant tillage treatment differences within soil fraction: *** = $p < 0.001$, ** = $p < 0.01$, * = $p < 0.05$. Speckled bars represent occluded fractions.

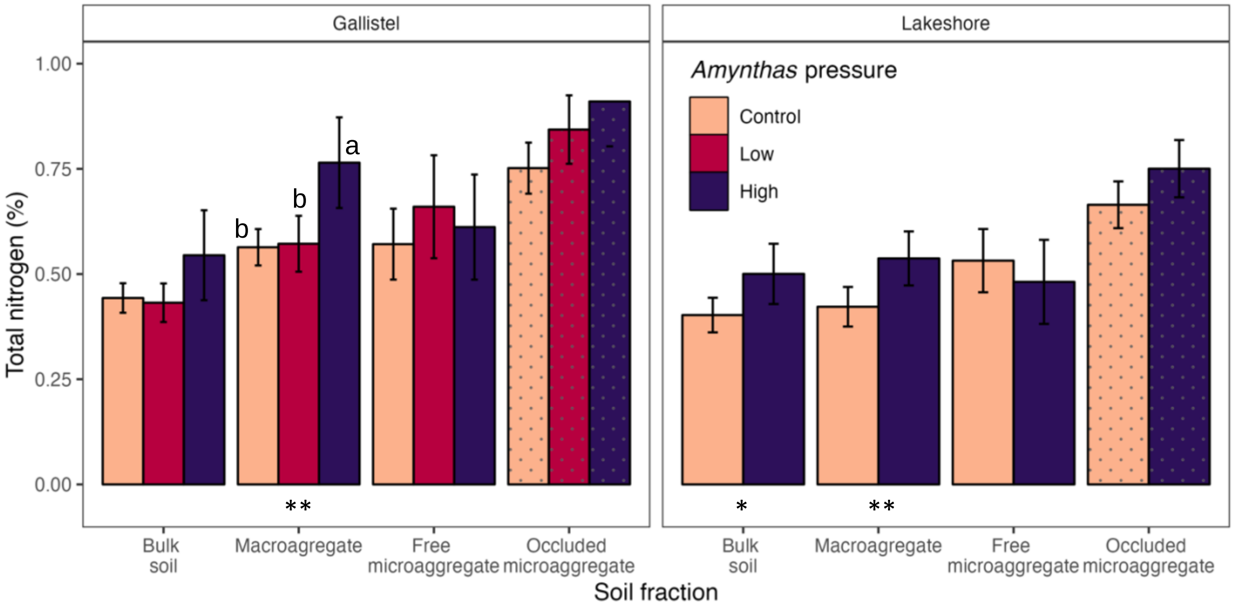

**Figure S4**. Nitrogen concentration (total N, %), by soil fraction and *Amynthas* pressure treatment. Soil fractions are as follows: Bulk soil = sieved field-moist soil < 2000 µm; Macroaggregate = macroaggregate fraction, 250–2000 µm; Free microaggregate = microaggregate fraction from bulk soil, 53–250 µm; Occluded microaggregate = microaggregate fraction occluded within macroaggregate fraction, 53–250 µm. Error bars represent ± 1.96 SE. Asterisks indicate significant tillage treatment differences within soil fraction: *** = $p < 0.001$, ** = $p < 0.01$, * = $p < 0.05$. Speckled bars represent occluded fractions.

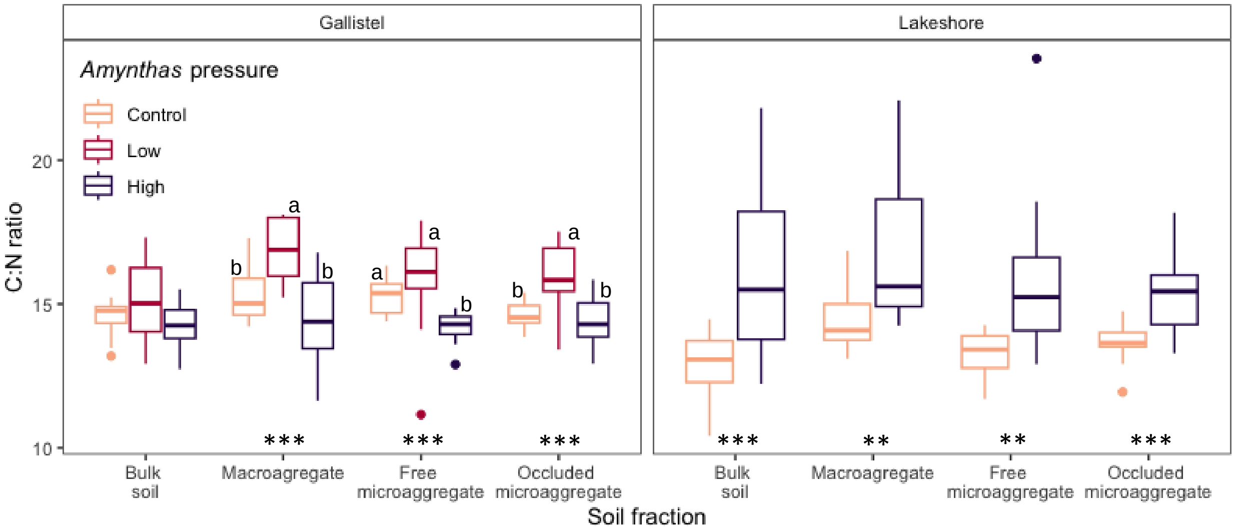

**Figure S5**. C:N ratio, by soil fraction and *Amynthas* pressure treatment. Soil fractions are as follows: Bulk soil = whole soil; Macroaggregate = macroaggregate fraction, 250–2000 µm; Free microaggregate = microaggregate fraction from bulk soil, 53–250 µm; Occluded microaggregate = microaggregate fraction occluded within macroaggregate fraction, 53–250 µm. Asterisks indicate significant tillage treatment differences, within soil fraction: *** = $p < 0.001$, ** = $p < 0.01$, * = $p < 0.05$. Different letters denote significant treatment differences within fraction at Gallistel.

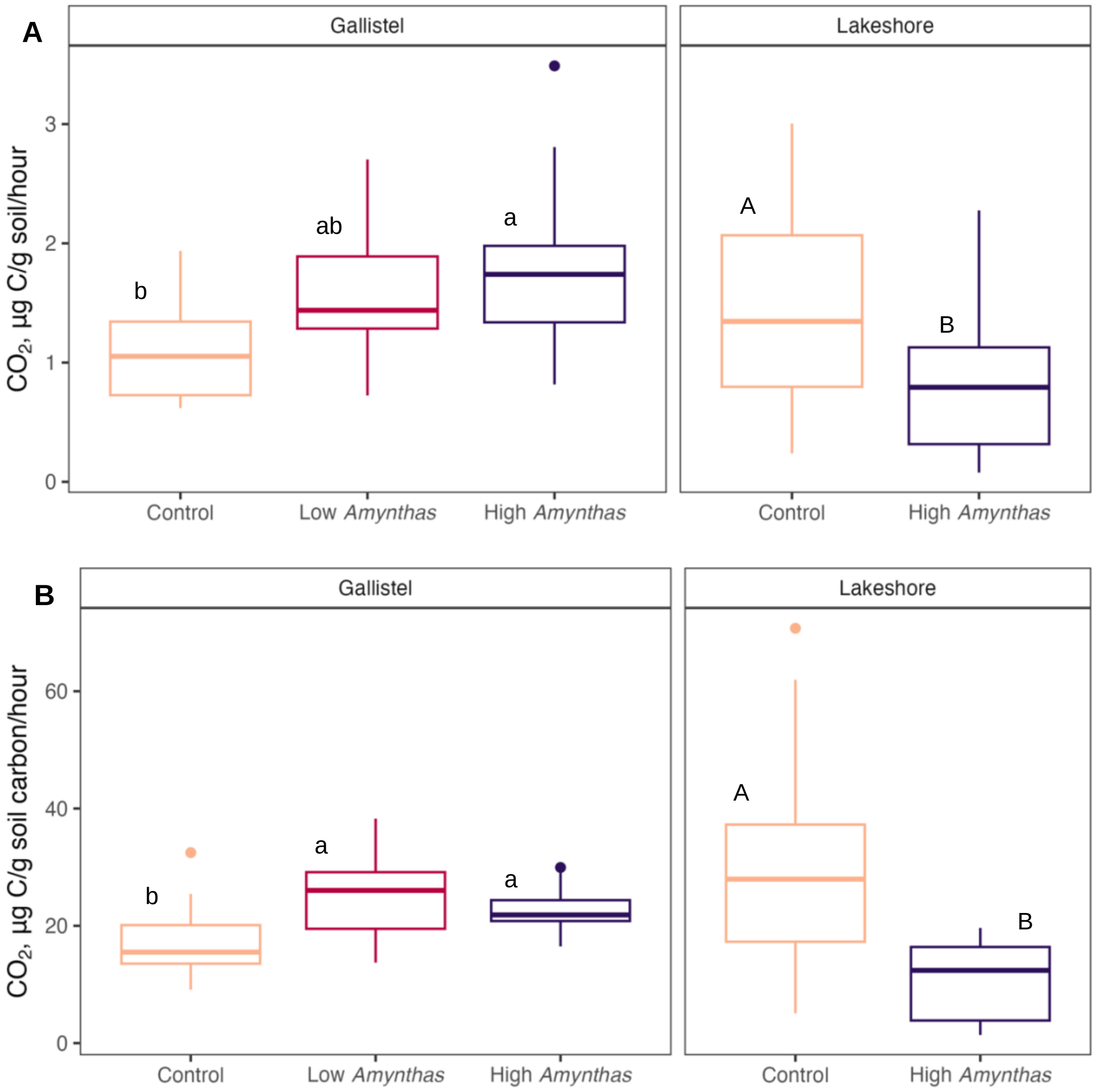

**Figure S6**. Soil respiration, by *Amynthas* pressure treatment on a per g soil basis (**A**) and on a per g soil carbon basis (**B**). $CO_2$ evolution was measured on field moist bulk soil sieved at 2 mm using the MicroResp system (Campbell et al., 2003), a setup designed to capture $CO_2$ from small soil samples using a colorimetric indicator mounted on top of a 96-deep well plate. Different letters denote significant treatment differences within site.

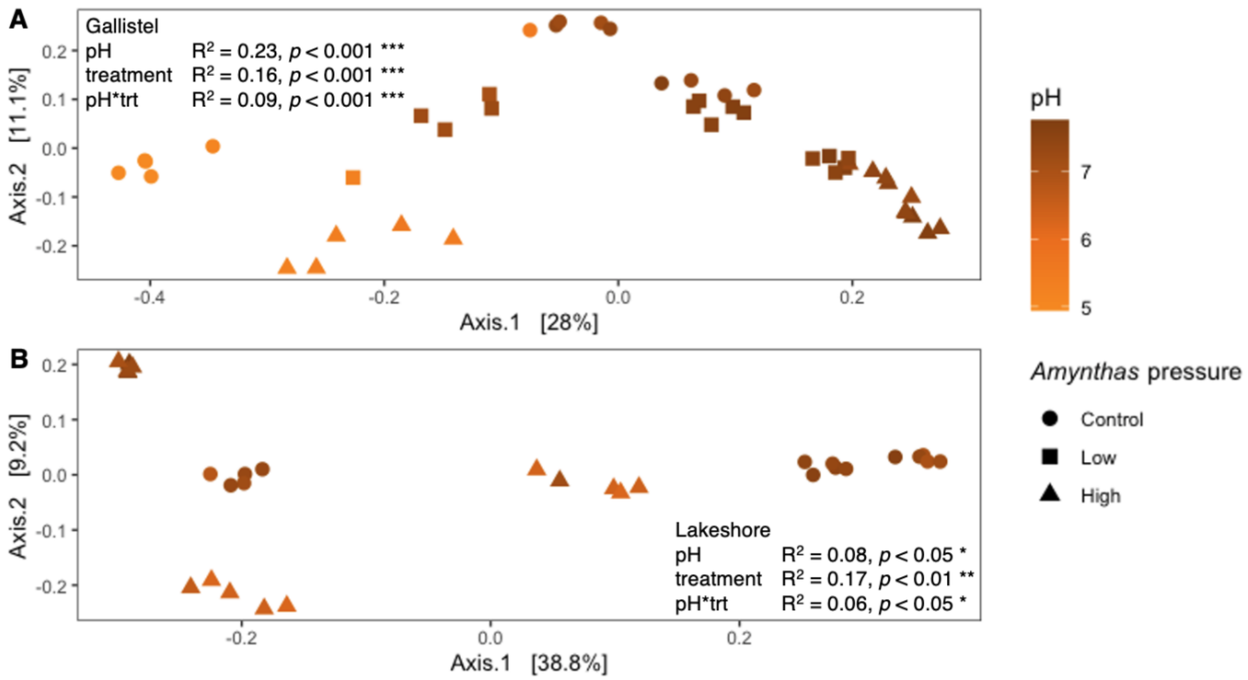

**Figure S7**. Principal coordinates analysis of Bray-Curtis dissimilarities of Hellinger-transformed community relative abundances, by soil pH and *Amynthas* pressure treatment for Gallistel (**A**) and Lakeshore (**B**) woodland sites in Madison, WI. Each point represents the community of one bulk soil sample. Displayed statistics are from PERMANOVA analysis.

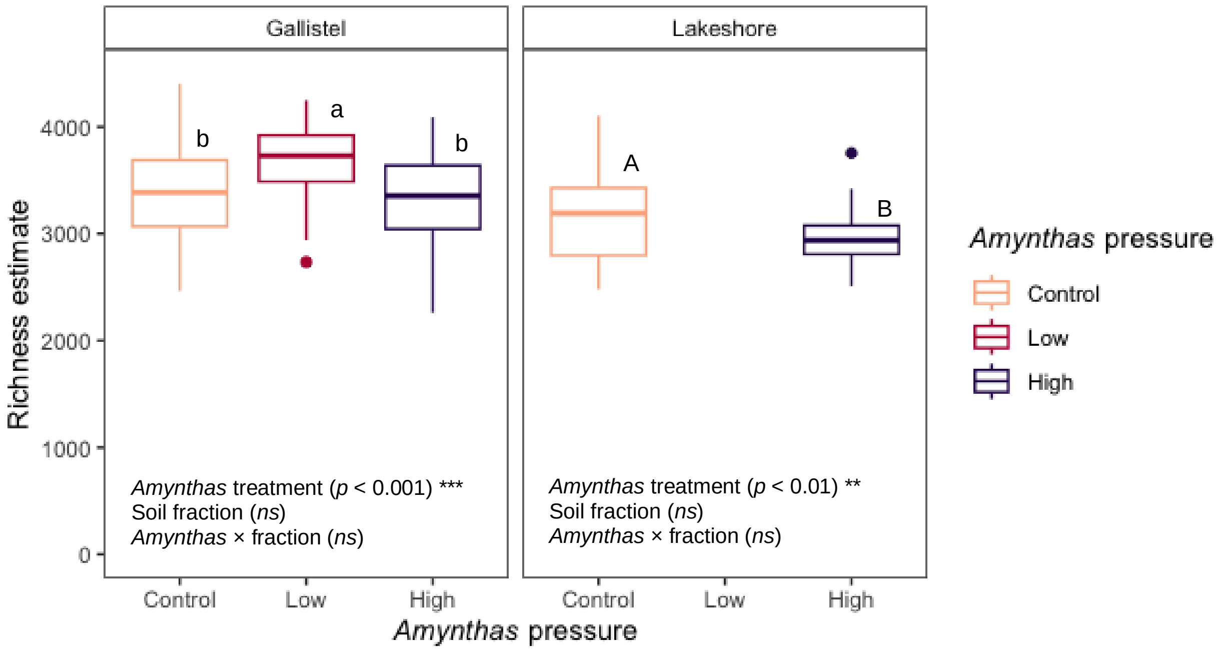

**Figure S8**. Bacterial OTU richness, by *Amynthas* treatment. Richness was estimated using the weighted linear regression model of OTU richness estimates, which weights observations based on variance, using *breakaway::betta* (Willis et al., 2017). Compact letter display, in which boxplots with the same letter are not significantly different, detail treatment differences within site.

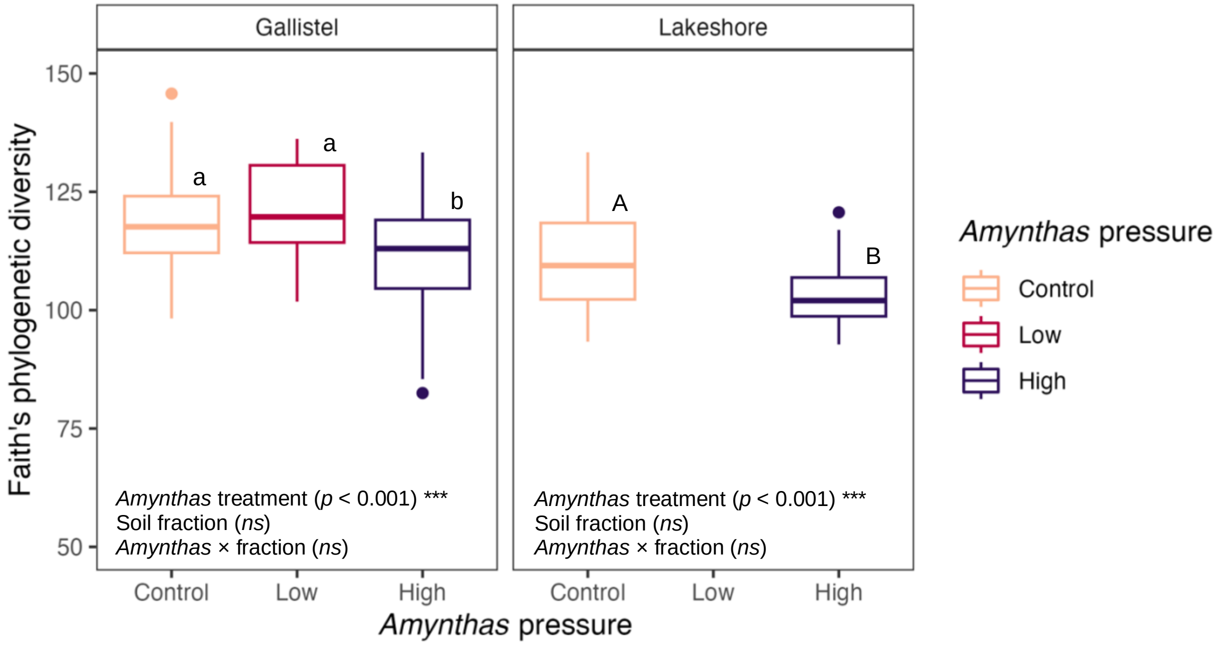

**Figure S9**. Faith's phylogenetic diversity, by *Amynthas* treatment. Asterisks indicate significant soil fraction difference: *** = $p < 0.001$, ** = $p < 0.01$, * = $p < 0.05$. Compact letter display, in which boxplots with the same letter are not significantly different, detail treatment differences within site.

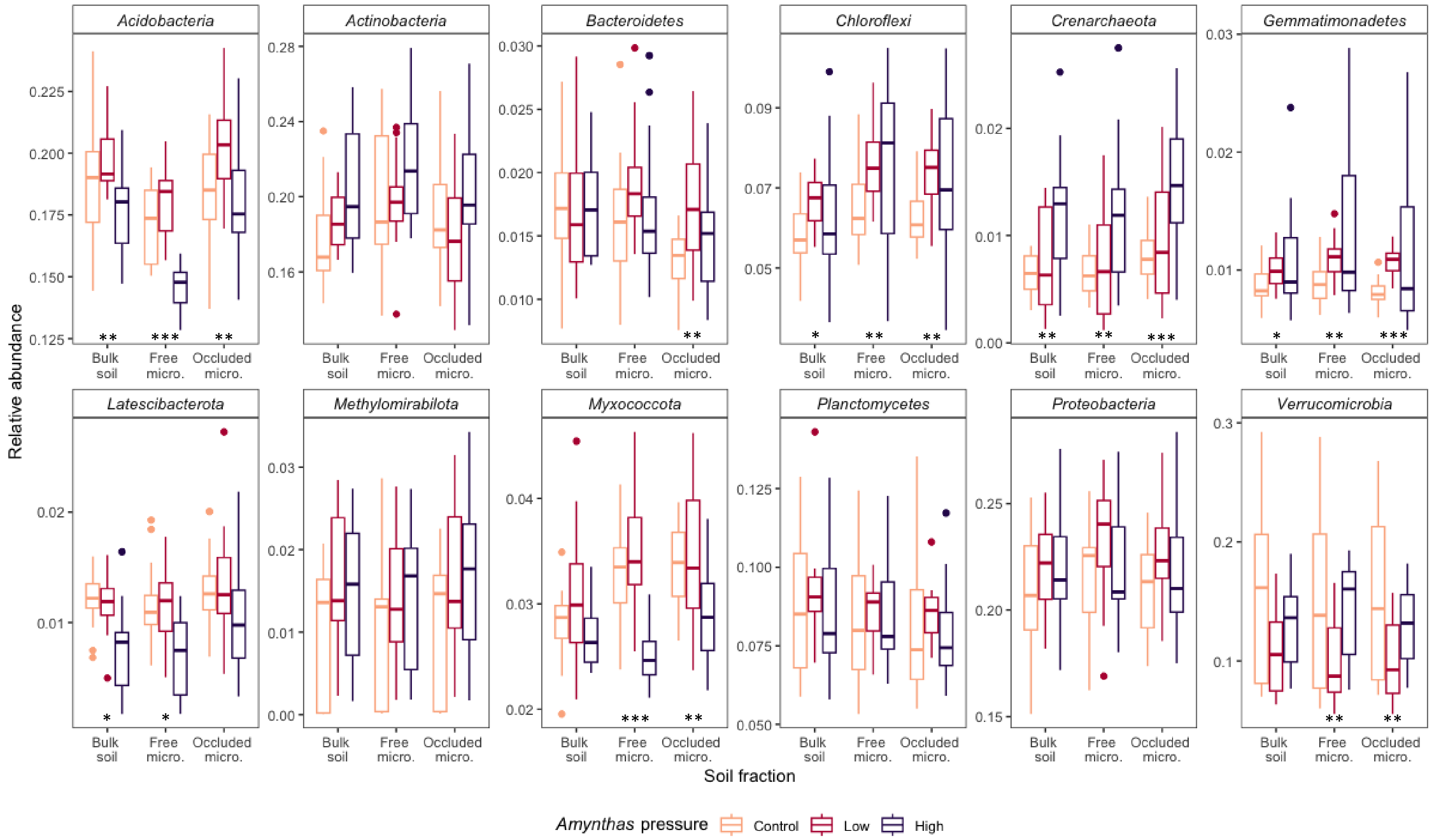

**Figure S10**. Relative abundances of representative phyla at Gallistel Woods site, Madison, WI. Bulk soil = whole soil; Free micro. = microaggregate fraction from bulk soil, 53–250 µm; Occluded micro. = microaggregate fraction occluded within macroaggregate fraction, 53–250 µm. Asterisks indicate significant treatment differences in relative abundances between tillage treatment, within fraction, determined using *corncob::differentialTest*, with taxa agglomerated at the phylum level: *** = $p < 0.001$, ** = $p < 0.01$, * = $p < 0.05$.

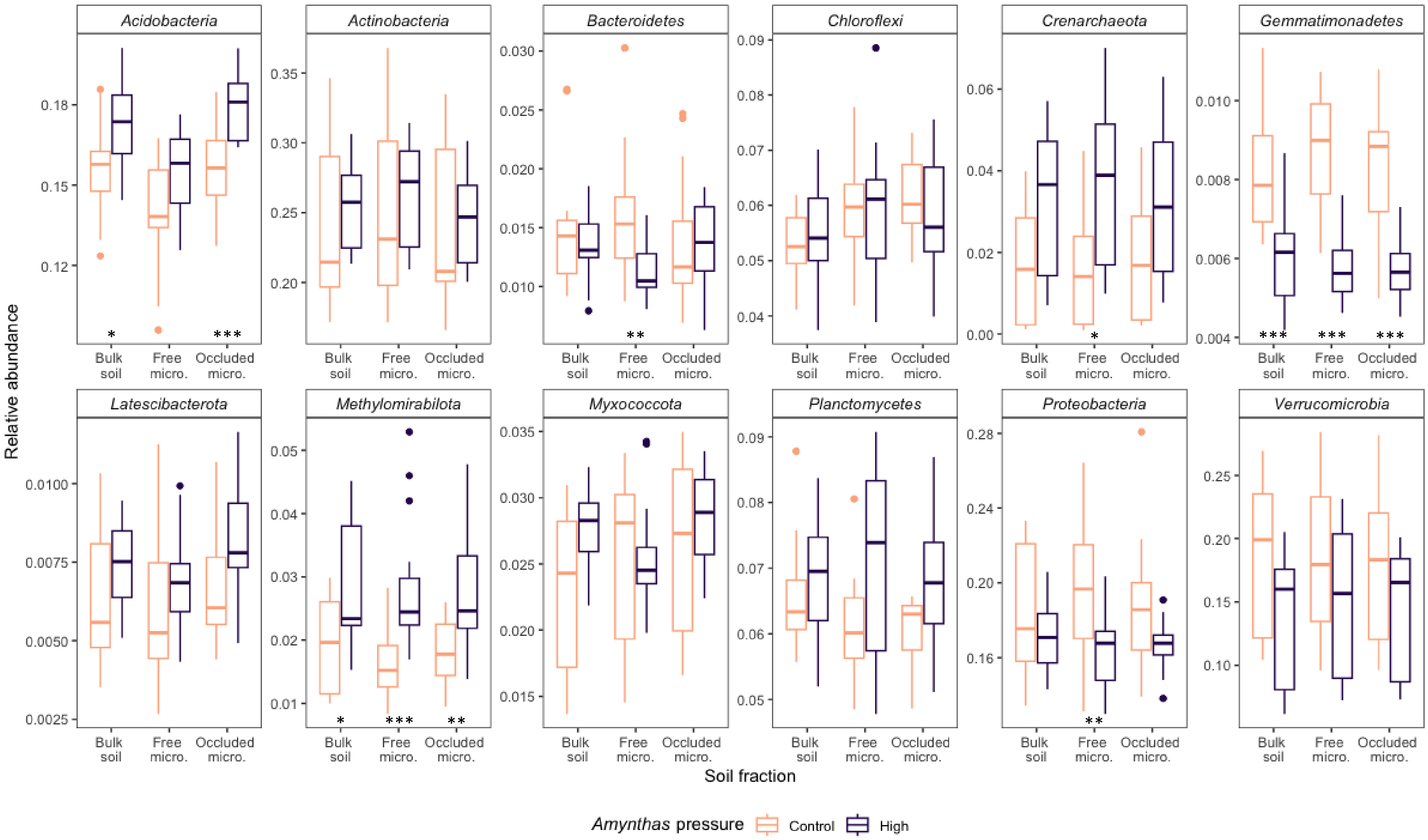

**Figure S11**. Relative abundances of representative phyla at Lakeshore Nature Preserve site in Madison, WI. Bulk soil = whole soil; Free micro. = microaggregate fraction from bulk soil, 53–250 µm; Occluded micro. = microaggregate fraction occluded within macroaggregate fraction, 53–250 µm. Asterisks indicate significant treatment differences in relative abundances between tillage treatment, within fraction, determined using *corncob::differentialTest*, with taxa agglomerated at the phylum level: *** = $p < 0.001$, ** = $p < 0.01$, * = $p < 0.05$.

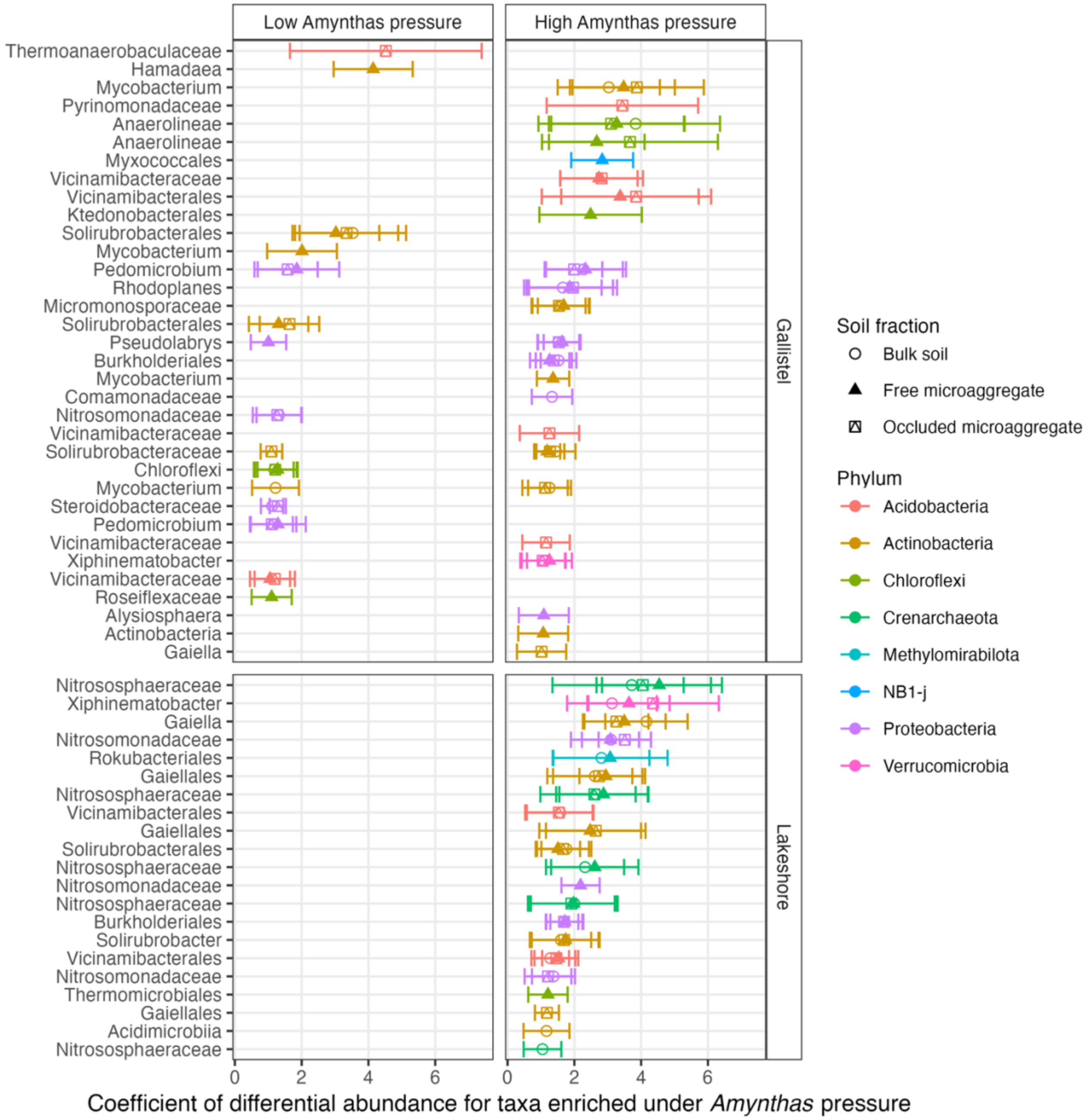

**Figure S12**. Taxa with positive differential abundance (enrichment) under low and high *Amynthas* pressure for each site. The x-axis is the coefficient of differential abundance, in this case (showing enrichment), demonstrating an increase in relative abundances of taxa under tillage as compared to no-tillage, with only coefficients > 1.0 plotted. Each point represents a single OTU, colored by phylum, and labelled on the y-axis with the finest available taxonomy; there may be more than one OTU definable to the same name and level of taxonomy. Error bars represent ± 1.96 SE

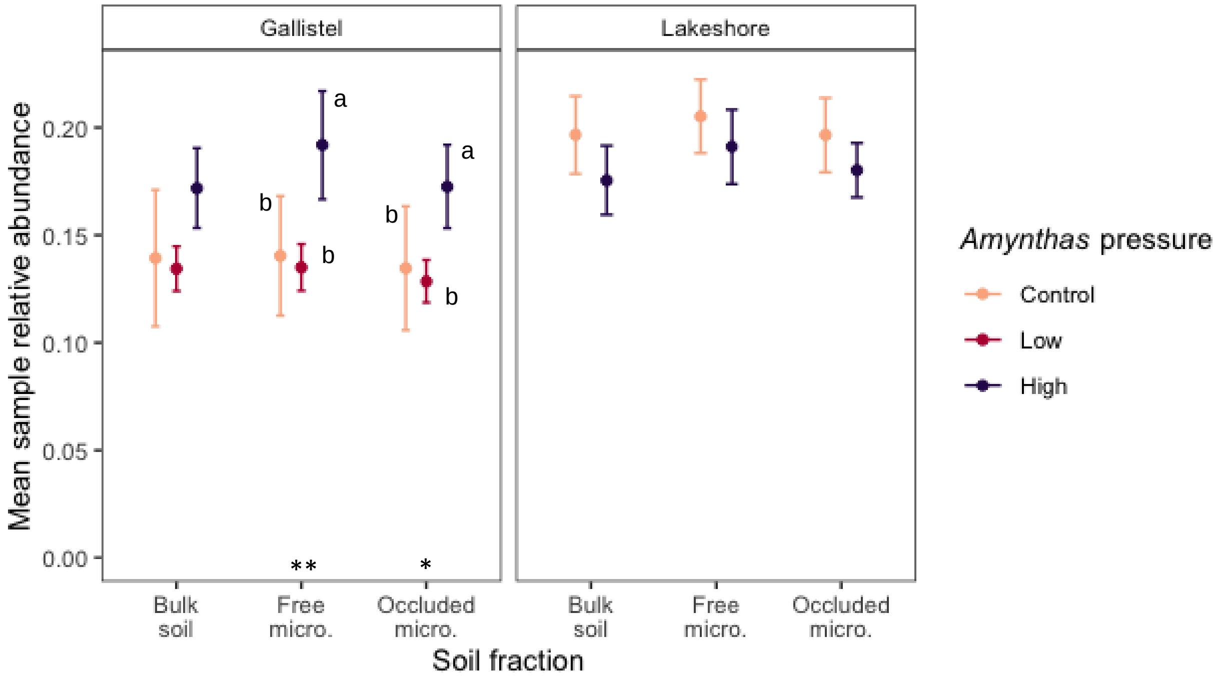

**Figure S13**. Mean relative abundance of the predominant worm casting taxa (taxa that comprise 75% of the worm casting relative abundance, by site) in soil. Bulk soil = whole soil; Free microagg. = microaggregate fraction from bulk soil, 53–250 µm; Occluded microagg. = microaggregate fraction occluded within macroaggregate fraction, 53–250 µm. Error bars represent 95% confidence interval (± 1.96 * standard error). Different letters, within soil fraction, designate statistically significant treatment differences ($p < 0.05$).

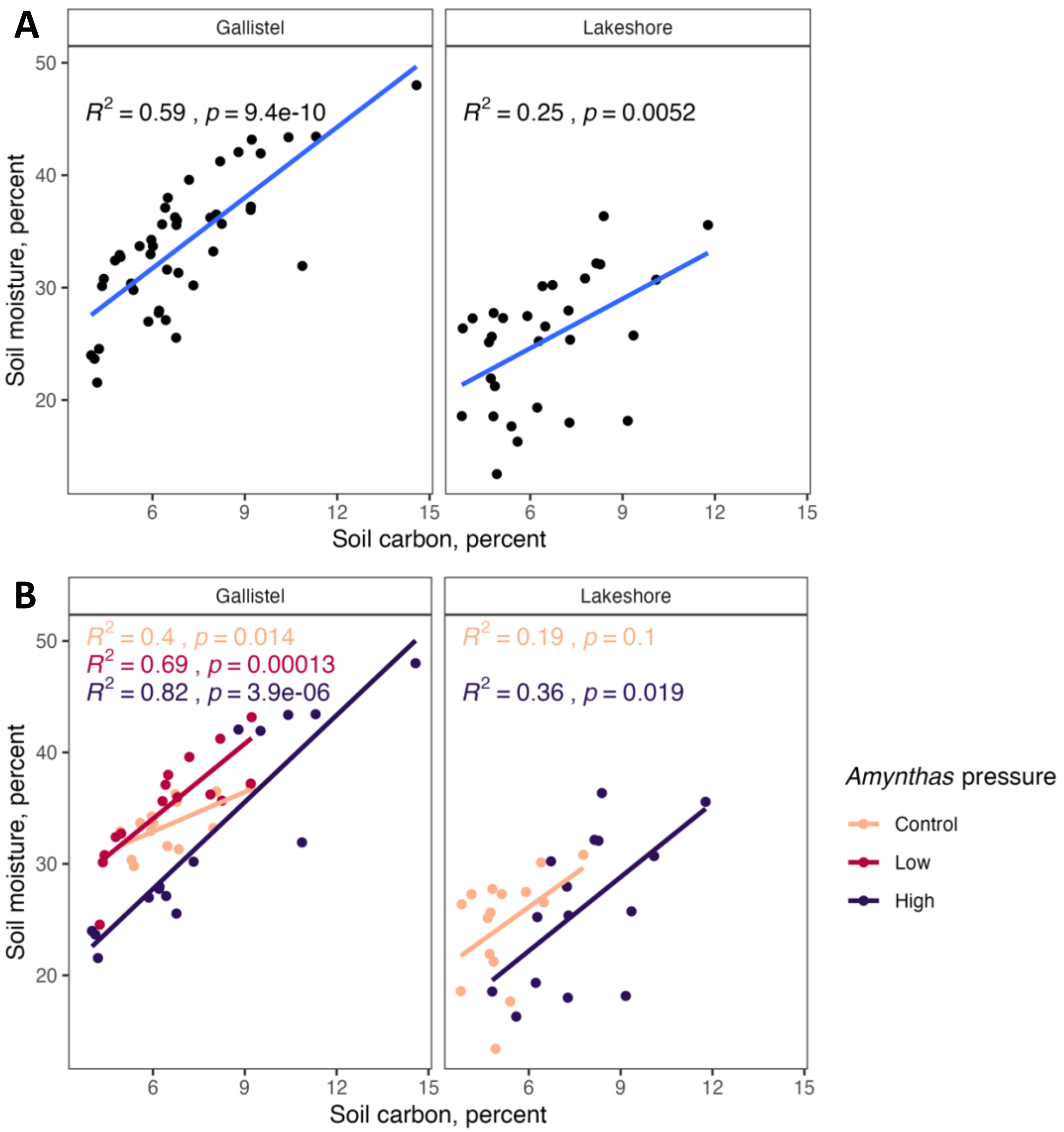

**Figure S14**. Relationship between soil moisture and soil carbon concentration of freshly collected, sieved, bulk soil, separated by site (**A**), and by *Amynthas* pressure treatment and site (**B**).

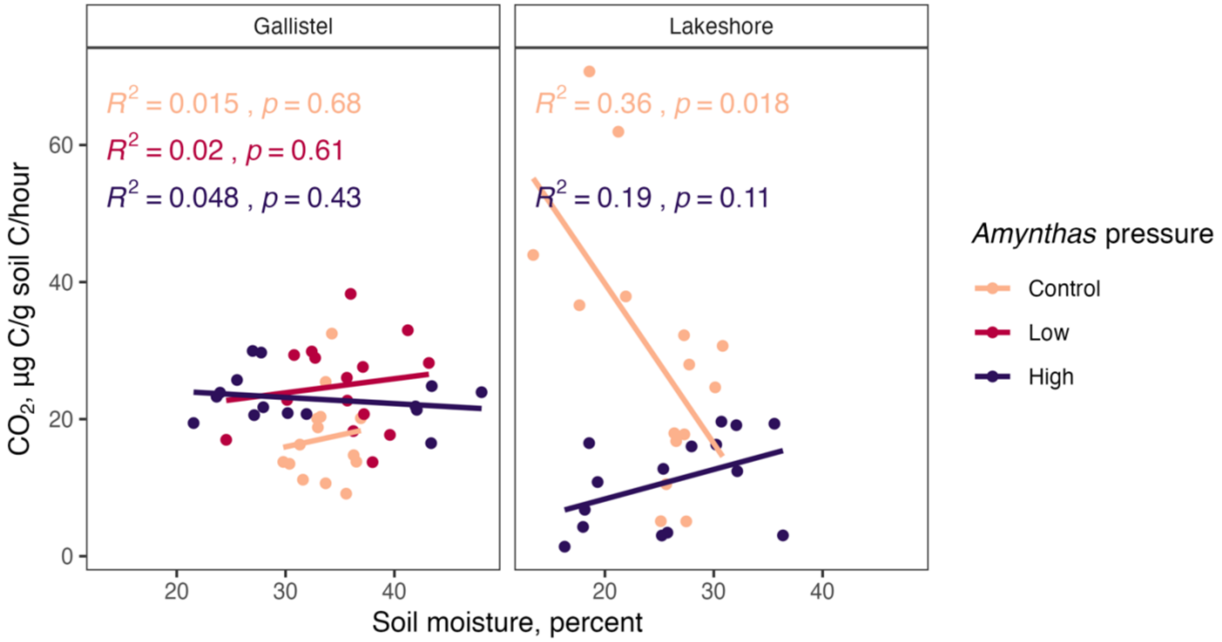

**Figure S15**. Relationship between soil respiration and soil moisture of freshly collected, sieved, bulk soil, separated by site and *Amynthas* pressure treatment. Respiration is reported here on a per unit of soil carbon basis, thus controlling for the effect of carbon content on respiration.
